# Redox heterogeneity as an engine of biodiversity: A quantitative murburn formalism for micro-oxic ecosystems

**DOI:** 10.64898/2026.08.19.745798

**Authors:** Kelath Murali Manoj, Abhinav Parashar

## Abstract

Biodiversity frequently peaks in fluctuating micro-oxic environments such as marine oxygen minimum zone interfaces, rhizospheric aggregates, sediments, microbial mats, and gut mucus layers. Yet, classical ecological theories do not adequately explain why intermediate oxygen tensions repeatedly favor coexistence and diversification. Herein, we propose a murburn ecological formalism wherein oxygen acts not merely as a metabolic substrate but as a generator of dynamic redox heterogeneity through partial reduction and diffusible reactive species (DRS) and redox-intermediates formation. Integrating empirical observations from marine, gut, soil, and aquatic-interface ecosystems with a reaction–diffusion framework, we show that intermediate oxygen tensions naturally maximize radical-field heterogeneity and produce dynamically shifting fitness landscapes. Numerical simulations demonstrate spontaneous coexistence, biodiversity maxima within micro-oxic zones, localized diversification, and coexistence stabilization without externally imposed niche partitioning. Additional simulations suggest that aquatic macrofauna indirectly enhance biodiversity by restructuring oxygen gradients and generating ecosystem-scale diffusional redox architectures (ESDRA). The framework proposes that fluctuating redox interfaces function as potential ecological zones of elevated adaptive turnover across biological scales.

## 1. Introduction

The origin, maintenance, and spatial organization of biodiversity remain among the most fundamental unresolved problems in ecology and evolutionary biology. Classical ecological theories explain coexistence primarily through mechanisms such as resource partitioning, trophic specialization, environmental filtering, dispersal limitation, stochastic drift, and adaptive evolution (Levin, 1992; Murray, 2002). While these approaches have yielded powerful conceptual and mathematical frameworks, they generally treat metabolism and environmental chemistry as passive background conditions rather than active dynamical generators of ecological structure.

Oxygen, in particular, is usually represented as a monotonic energetic resource. Conventional ecological and physiological models typically assume that increasing oxygen availability enhances aerobic viability until toxicity thresholds are reached, whereas oxygen depletion progressively restricts metabolic capacity. Such frameworks are sufficient for describing broad energetic trends, yet they fail to explain a striking and repeatedly observed ecological pattern: biodiversity frequently peaks not in fully oxic or fully anoxic environments, but in fluctuating micro-oxic systems where oxygen concentrations remain low but non-zero (Diaz & Rosenberg, 2008; Wright et al., 2012).

This phenomenon is observed across remarkably diverse ecological contexts, including: marine oxygen minimum zone (OMZ) interfaces, estuarine transition layers, microbial mats, rhizospheric soil aggregates, wetland sediments, biofilms, coral-associated microbiomes, and gut mucus interfaces (Falkowski et al., 2008; Albenberg et al., 2014; Kuzyakov & Blagodatskaya, 2015). These environments share several physicochemical characteristics:

1. steep oxygen gradients,
2. fluctuating diffusion barriers,
3. coexistence of multiple oxidation states,
4. elevated radical and redox activity,
5. and strong spatiotemporal heterogeneity.

Despite the robustness of these empirical observations, conventional ecological frameworks do not provide a mechanistically satisfying explanation for why intermediate oxygen tensions repeatedly favor coexistence, diversification, and rapid adaptive turnover.

Similarly, classical redox biology also faces conceptual limitations in such environments. Reactive oxygen species (ROS), reactive nitrogen species (RNS), halogen reactive intermediates, and related diffusible intermediates are generally interpreted as toxic byproducts or secondary signaling modifiers superimposed upon otherwise deterministic metabolic pathways. However, growing evidence from microbiology, systems biology, environmental physiology, geochemistry, and immunology indicates that diffusible reactive species (DRS; in the current context, these can also connote redox-active intermediates) participate directly in metabolism, signaling, extracellular coupling, environmental transformation, and ecological interaction (Apel & Hirt, 2004; Falkowski et al., 2008; Diaz et al., 2013; Hansel et al., 2019; Hansel & Diaz, 2021)

Extracellular superoxide generation by marine microbes, peroxide cycling in phytoplankton communities, ROS-mediated soil polymer degradation, epithelial ROS generation in gut interfaces, and metal-catalyzed redox cycling in sediments all demonstrate that reactive intermediates are not rare pathological artifacts but constitutive components of natural ecosystems (Diaz et al., 2013; Hansel et al., 2019). Furthermore, many ecological transitions (including algal blooms, dysbiosis, hypoxia-associated microbial restructuring, and soil succession) display strongly nonlinear threshold behavior suggestive of chemically mediated feedback systems rather than simple deterministic resource competition.

Murburn concept provides a physicochemical framework within which these observations become mechanistically coherent. Murburn (“*mured burning*”) proposes that DRS or redox intermediates generated through partial reduction and stochastic redox interactions are not merely secondary consequences of metabolism but integral mediators of bioenergetics, signaling, adaptation, and environmental coupling (Manoj, 2025). Rather than viewing biological systems as tightly wired deterministic electron-transfer assemblies, murburn logic emphasizes stochastic redox dynamics, diffusional interactions, distributed reaction zones (“murzones”), and non-equilibrium radical-mediated chemistry.

When extended to ecology, murburn principles imply that oxygen functions not solely as a terminal electron acceptor or energetic substrate, but as a generator of dynamic redox heterogeneity. In fluctuating micro-oxic systems, partial oxygen reduction produces transient spectra of diffusible reactive intermediates whose concentrations depend upon oxygen diffusion, substrate availability, environmental structure, transition-metal chemistry, and organismal activity.

Under such conditions, ecological fitness becomes intrinsically redox-dependent. Organisms differing in radical tolerance, radical utilization, antioxidant buffering, diffusional positioning, and metabolic flexibility may each gain transient advantages under different local conditions. Because the redox environment is itself dynamically modified by biological activity, ecological feedback becomes nonlinear and spatially heterogeneous. Consequently, competitive exclusion is suppressed, coexistence becomes chemically scaffolded, and fluctuating micro-oxic environments function as potential zones of elevated adaptive turnover (serving as sorts of evolutionary incubators!).

The central proposition advanced herein is therefore that intermediate oxygen tensions maximize stochastic redox heterogeneity and DRS flux, thereby generating dynamically shifting fitness landscapes that stabilize coexistence and accelerate diversification.

To formalize this proposition, we develop a mathematically minimal murburn ecological framework based on coupled reaction–diffusion equations linking oxygen gradients, radical-field dynamics, and redox-dependent organismal fitness. We then relate this formalism to ecological observations from marine oxygen minimum zones, gut mucus microinterfaces, and rhizospheric soil aggregates.

The resulting framework proposes that biodiversity is fundamentally coupled to the spatial and temporal organization of redox heterogeneity, with micro-oxic niches acting as chemically dynamic engines of ecological complexity and evolutionary innovation.

## 2. Empirical ecological foundation

### 2.1. Biodiversity maxima in fluctuating oxygen environments

Across marine, terrestrial, and host-associated ecosystems, biodiversity frequently peaks in environments characterized by fluctuating oxygen availability rather than fully oxic or fully anoxic conditions (Diaz & Rosenberg, 2008; Wright et al., 2012). Such observations recur despite profound differences in phylogeny, spatial scale, nutrient architecture, and environmental context, suggesting the existence of a general organizing principle linked to redox dynamics.

Micro-oxic environments share several physicochemical characteristics:

1. oxygen concentrations remain low but non-zero,
2. diffusion is spatially constrained,
3. multiple oxidation states coexist simultaneously,
4. partial oxygen reduction is favored,
5. reactive intermediates accumulate transiently,
6. and redox conditions fluctuate across micron-to-centimeter scales.

These conditions naturally promote chemically heterogeneous environments in which no single metabolic strategy remains universally optimal.

Under classical ecological logic, coexistence generally requires stable niche partitioning or tradeoffs among fixed resource-utilization strategies. However, fluctuating micro-oxic systems do not behave as chemically static environments. Instead, environmental chemistry itself becomes dynamic, spatially heterogeneous, and biologically self-modifying. Murburn theory proposes that these fluctuating radical fields are themselves ecological structuring agents.

### 2.2. Marine oxygen minimum zones (OMZs)

Marine oxygen minimum zones constitute among the most chemically dynamic ecosystems on Earth. OMZs arise where biological oxygen consumption exceeds physical replenishment, producing stratified water columns containing steep transitions between oxic surface waters and near-anoxic deeper regions (Wright et al., 2012).

Major OMZ systems occur in: the Eastern Tropical Pacific, the Arabian Sea, the Benguela upwelling system, and semi-enclosed basins such as Saanich Inlet. Importantly, biodiversity and metabolic complexity often peak near OMZ transition boundaries rather than within fully oxygenated or fully anoxic waters (Wright et al., 2012; Canfield et al., 2010). These interfaces support dense and metabolically diverse microbial consortia involved simultaneously in: aerobic respiration, denitrification, sulfur oxidation, methane oxidation, nitrate reduction, and anaerobic ammonium oxidation (anammox) (Ward, 2013). Such coexistence is difficult to reconcile using strictly compartmentalized respiratory logic because classical deterministic frameworks predict sharper exclusion among competing respiratory modes. Instead, OMZ interfaces display: overlapping energetic pathways, metabolic redundancy, rapid ecological turnover, and extensive functional overlap. Marine OMZ systems are also characterized by intense redox cycling involving: dissolved oxygen reactive intermediates, sulfur intermediates, metal-ion catalysis, and photochemically generated ROS. Extracellular superoxide production by marine bacteria and phytoplankton has now been experimentally demonstrated across multiple oceanic systems (Diaz et al., 2013; Hansel et al., 2019). Hydrogen peroxide gradients correlate strongly with biological productivity, photochemical activation, and microbial turnover (Hansel & Diaz, 2021).

From a murburn perspective, OMZ interfaces behave as distributed stochastic redox reactors. Low but non-zero oxygen availability promotes partial oxygen reduction and transient ROS generation, while dissolved metals, sulfides, and organic substrates participate in diffusional radical cycling. These interactions generate spatially dynamic redox mosaics in which distinct physiological strategies remain locally advantageous under continuously shifting conditions. Importantly, marine organisms themselves actively reshape local redox fields through: oxygen consumption, extracellular metabolite release, peroxide generation, sulfur cycling, and radical scavenging. Ecological structure therefore emerges from reciprocal coupling between organisms and chemically mediated environmental heterogeneity. This interpretation is strongly compatible with empirical observations showing that OMZ transition regions often exhibit maximal microbial diversity, functional redundancy, and rapid adaptive turnover (Wright et al., 2012).

### 2.3. Gut mucus micro-oxic interfaces

The gastrointestinal tract provides another striking example of biodiversity concentration within micro-oxic interfaces. Although the intestinal lumen is predominantly anaerobic, epithelial surfaces remain relatively oxygenated because of vascular oxygen supply. Between these regions exists a steep oxygen gradient within the mucus layer, generating a highly dynamic micro-oxic interface (Albenberg et al., 2014; Zheng et al., 2015). This interface supports coexistence among: obligate anaerobes, facultative anaerobes, aerotolerant organisms, and microaerophiles. Importantly, epithelial and immune cells continuously generate reactive oxygen and nitrogen intermediates, while microbial populations simultaneously produce, utilize, scavenge, and buffer these species.

Experimental studies have shown that oxygen gradients strongly influence spatial partitioning of gut microbial communities (Albenberg et al., 2014). Inflammatory conditions further demonstrate the ecological importance of redox modulation. Small perturbations in epithelial oxygen leakage or ROS balance can induce large-scale microbiome restructuring, favoring taxa with greater oxidative tolerance while suppressing strictly anaerobic competitors. These ecological transitions frequently occur abruptly and nonlinearly, suggesting threshold-like chemically mediated dynamics rather than gradual resource replacement alone (Albenberg et al., 2014; Donaldson et al., 2016).

Under murburn assumptions, the gut mucus interface behaves as a dynamic redox field in which diffusible reactive intermediates continuously reshape local viability landscapes. Spatially heterogeneous radical fluxes generate chemically differentiated micro-niches permitting coexistence among multiple metabolic strategies. Because organisms simultaneously modify local oxygen consumption and redox buffering capacity, the ecosystem becomes self-organizing through reciprocal chemical feedback. The result is a fluctuating coexistence system rather than a stable deterministic hierarchy. Gut biodiversity therefore emerges not merely from trophic partitioning or host selection but from chemically mediated redox heterogeneity operating across microscopic spatial scales.

This interpretation aligns strongly with observations that: (a) moderate oxidative gradients support maximal microbial complexity, and (b) whereas severe inflammation or complete oxygen depletion both reduce biodiversity (Zheng et al., 2015).

### 2.4. Rhizosphere and soil aggregate systems

Soil ecosystems exhibit perhaps the most spatially fragmented oxygen architecture in nature. Air-filled pores, water-saturated microdomains, fungal hyphae, decomposing organic aggregates, and plant-root interfaces together generate steep and rapidly fluctuating oxygen gradients across micron-scale distances (Vos et al., 2013; Kuzyakov & Blagodatskaya, 2015). Microbial diversity is especially elevated within rhizospheric and aggregate-associated micro-oxic regions where oxic and anoxic domains intersect. These environments support extraordinary metabolic diversity involving: aerobic respiration, fermentation, denitrification, methanogenesis, sulfur cycling, extracellular oxidative chemistry, and metal-dependent redox transformations. Transition metals such as iron and manganese further contribute to dynamic radical generation through Fenton-type chemistry and related redox cycling processes. Simultaneously, lignin degradation, humic oxidation, and extracellular polymer decomposition involve diffusible oxidative intermediates extending beyond rigid enzymatic active sites.

The rhizosphere also contains highly heterogeneous nutrient distributions, fluctuating hydration conditions, and intense microbial competition, all of which amplify redox variability.

Classical deterministic metabolic frameworks inadequately explain how so many partially overlapping physiological strategies coexist within such confined spatial domains. Under murburn assumptions, soil aggregates behave as distributed stochastic redox reactors. Partial oxygenation continuously generates reactive intermediates whose concentrations fluctuate with: hydration state, diffusional barriers, substrate availability, metal-ion chemistry, and microbial activity.

These chemically heterogeneous conditions prevent long-term dominance by any single energetic strategy. Instead, shifting redox microdomains favor dynamic coexistence among organisms possessing differing: oxidative tolerances, buffering systems, diffusional positioning, and metabolic flexibility. The rhizosphere therefore becomes an ecological and evolutionary mosaic structured by redox-field heterogeneity rather than by static metabolic compartmentalization alone.

### 2.5. Aquatic macrofauna and ionically mediated redox ecology

Aquatic ecosystems provide uniquely favorable physicochemical conditions for the emergence and spatial propagation of murburn-compatible redox dynamics. Unlike terrestrial systems, where gaseous diffusion dominates and reactive intermediates are often rapidly dispersed or quenched, aquatic environments are fundamentally water-mediated reaction spaces in which diffusion, ionic interactions, interfacial chemistry, and radical stabilization become ecologically significant across multiple spatial scales. Consequently, water-rich ionic environments transform localized radical chemistry into ESDRA.

This distinction is especially important because aquatic ecosystems contain: persistent dissolved oxygen gradients, extensive ionic conductivity, transition-metal redox cycling, halide-rich chemistries, photochemically active interfaces, and highly structured biological boundary layers. Together, these properties amplify the ecological consequences of murburn-like stochastic redox processes.

Under classical ecological interpretations, water primarily functions as a transport medium or solvent background. In contrast, the murburn framework suggests that aquatic environments actively shape ecological organization by modulating radical diffusion, persistence, stabilization, and interfacial coupling.

#### 2.5.1. Water abundance changes diffusion-limited redox behavior

Water-rich systems fundamentally alter diffusion-limited redox dynamics because oxygen diffusion in aqueous environments is substantially slower than in air. Consequently, aquatic systems readily develop persistent oxygen gradients across relatively short spatial scales (Revsbech & Jørgensen, 1986). Such gradients occur in: marine snow particles, mucus layers, sediment interfaces, algal biofilms, coral surfaces, gill structures, and rhizospheric aggregates.

These micro-oxic environments are particularly favorable for partial oxygen reduction and transient DRS accumulation. Importantly, aqueous environments also prolong the ecological significance of diffusible reactive intermediates. Superoxide, peroxide, hydroxyl radicals, and related species can participate in relay chemistry, hydration-shell interactions, and transition-metal-mediated propagation processes that extend reactive influence beyond localized enzymatic sites (Diaz et al., 2013; Hansel et al., 2019).

Under murburn assumptions, water therefore functions not merely as a passive solvent but as a diffusional and dielectric matrix capable of spatially organizing stochastic redox interactions. Because oxygen penetration, radical generation, and radical quenching all become diffusion-limited processes in aqueous systems, aquatic ecosystems naturally generate dynamic redox mosaics.

This interpretation aligns strongly with experimental observations demonstrating steep oxygen microgradients and ROS-associated ecological restructuring within aquatic microbial communities, biofilms, and sediment interfaces (Wright et al., 2012; Falkowski et al., 2008). Moreover, because water simultaneously stabilizes ionic interactions and constrains oxygen replenishment, fluctuating micro-oxic domains become persistent rather than transient ecological structures. Such conditions strongly favor coexistence among organisms possessing differing oxidative tolerances, diffusional positioning, and radical-buffering capacities. Thus, water abundance itself amplifies the ecological consequences of murburn-like dynamics by converting localized redox events into spatially extended ecological fields.

#### 2.5.2. Halides are extremely important

Marine systems differ profoundly from most terrestrial environments because seawater is highly ionic and halide-rich. Chloride concentrations in seawater approach approximately 0.5 M, while bromide and iodide are also present at ecologically meaningful concentrations. These ions are not chemically inert background species. Instead, they participate directly in oxidative and radical-mediated chemistry through haloperoxidase activity, halogenation reactions, and diffusible halogen-radical formation (La Barre et al., 2010).

Marine algae, cyanobacteria, sponges, mollusks, and associated microbiomes produce large numbers of halogenated metabolites and haloperoxidase enzymes capable of generating: hypochlorous/hypobromous intermediates, halogen reactive intermediates, singlet oxygen species, and oxidized halometabolites. Such chemistry substantially expands the ecological significance of DRS beyond oxygen-centered reactive intermediates alone. Halides are especially important because they:

1. participate in diffusible oxidative relay systems,
2. modify radical persistence,
3. alter interfacial electrochemistry,
4. influence microbial competition,
5. and contribute to extracellular oxidative structuring.

Halogenated oxidative chemistry has already been implicated in: marine biofilm regulation, antifouling defenses, microbial competition, coral-associated chemistry, and algal ecological interactions. From a murburn perspective, seawater therefore acts as a chemically conductive radical-buffering matrix in which halide-dependent redox processes amplify ecological heterogeneity. This interpretation is strongly compatible with observations that marine ecosystems often exhibit exceptionally complex chemical ecologies involving extracellular oxidants, diffusible metabolites, and oxidative signaling networks (Falkowski et al., 2008).

Importantly, the present framework does not propose that halogen reactive intermediates universally dominate aquatic ecology. Rather, it suggests that halide-rich ionic environments amplify and spatially extend murburn-compatible redox dynamics.

#### 2.5.3. Marine interfaces are radical hotspots

Many marine ecological interfaces behave as natural radical hotspots because they combine: fluctuating oxygen tension, high ionic strength, dense biological activity, transition-metal cycling, photochemical activation, and diffusion-limited transport. Examples include: coral mucus layers, gill surfaces, estuarine boundaries, marine snow aggregates, sediment-water interfaces, phytoplankton blooms, and biofouling films.

These systems are characterized by steep oxygen gradients and intense extracellular chemistry. Simultaneously, biological metabolism continuously modifies local redox conditions through oxygen consumption, sulfur cycling, peroxide release, and extracellular metabolite production.

Marine snow particles are particularly illustrative. These suspended organic aggregates contain highly structured micro-oxic interiors surrounded by relatively oxygen-rich waters. Experimental studies demonstrate that such particles support extraordinarily diverse microbial consortia exhibiting overlapping respiratory and metabolic strategies. Under murburn assumptions, marine snow aggregates function as localized stochastic redox reactors in which diffusional oxygen limitation and radical-field heterogeneity stabilize coexistence (Ploug, 2001; Stewart & Franklin, 2008).

Similarly, coral mucus layers exhibit steep chemical gradients and ROS-associated ecological interactions involving both host tissues and associated microbial communities. Coral bleaching itself has been strongly linked to oxidative imbalance and redox collapse under thermal stress.

The murburn framework interprets these systems as examples of ESDRA wherein localized radical chemistry becomes spatially propagated through aqueous ionic environments.

#### 2.5.4. Macroscopic fauna as redox landscape engineers

Macroscopic aquatic organisms do not merely inhabit chemically heterogeneous environments; they actively generate and reshape them. Fish, mollusks, corals, crustaceans, echinoderms, annelids, and other aquatic fauna continuously modify:oxygen distributions, mucus-layer chemistry, ion gradients, peroxide fluxes, sulfur cycling, and microbial redox environments. For example: gill ventilation alters local oxygen penetration, mucus secretion modifies diffusion barriers, benthic burrowing restructures sediment oxygenation, filter feeding reshapes microbial distributions, and coral-associated mucus production generates chemically active interfacial layers. These processes create dynamic micro-oxic niches whose physicochemical properties are biologically regulated (Jones et al., 1994; Laland et al., 2016). Under murburn assumptions, such organisms therefore function as redox landscape engineers. Their activities continuously reshape local DRS fields and diffusional architectures, thereby influencing ecological coexistence, microbial succession, and nutrient cycling. Importantly, this interpretation does not require that macrofauna themselves operate primarily through murburn mechanisms internally. Rather, the ecological environments they generate amplify murburn-compatible stochastic redox interactions among associated microbial and mesoscopic communities. This distinction is critical because it preserves mechanistic restraint while still recognizing the ecosystem-scale consequences of organism-mediated redox structuring.

#### 2.5.5. Freshwater systems as comparative redox ecologies

Freshwater ecosystems provide an important comparative contrast to marine environments because they differ substantially in: ionic strength, halide abundance, conductivity, buffering capacity, and transition-metal availability. Consequently, freshwater systems may exhibit fundamentally different redox-field architectures despite sharing similar oxygen-gradient dynamics. Micro-oxic niches remain widespread in freshwater systems including: lake sediments, wetlands, stratified lakes, river biofilms, and freshwater rhizospheres. However, lower halide abundance may reduce the extent of halogen-mediated oxidative relay chemistry relative to marine systems. This distinction generates an important ecological prediction: marine ecosystems should exhibit more spatially extensive and chemically buffered diffusional redox architectures than freshwater systems.

At the same time, freshwater systems may exhibit greater sensitivity to localized oxygen collapse because reduced ionic buffering limits stabilization of fluctuating radical fields. Comparative investigation of marine versus freshwater redox ecology may therefore provide an experimentally tractable means of testing murburn ecological predictions. More broadly, the contrast between marine and freshwater environments highlights a central proposition of the present framework: the physicochemical properties of water-rich ionic environments profoundly influence how localized radical chemistry scales into ecosystem-level ecological organization.

## 3. Murburn quantitative-theoretical formulation

### 3.1. Conceptual basis

The murburn ecological formalism proposed herein is based on four core assumptions:

1. Oxygen gradients generate DRS through partial reduction processes.
2. Radical generation is maximal under intermediate oxygen availability.
3. Organismal fitness depends non-monotonically upon local redox intensity.
4. Organisms dynamically modify the same redox fields that govern their own viability.

Unlike classical ecological models, which typically assume fixed interaction coefficients and static environmental structure, the present framework treats ecological interactions as emergent consequences of spatially distributed redox chemistry. Because ecological systems are diffusion-limited, spatially heterogeneous, and chemically stochastic, reaction–diffusion formalism represents the minimal mathematically appropriate framework.

### 3.2. Core state variables

We define:

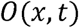

as the local oxygen concentration,

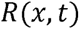

as the local concentration of DRS (ROS, RNS, halogen reactive intermediates, and related intermediates),

and

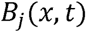

as the biomass density of species *j*.

The substrate pool is incorporated implicitly within radical-generation terms to maintain mathematical compactness.

### 3.3. Oxygen field dynamics

The oxygen field evolves according to:

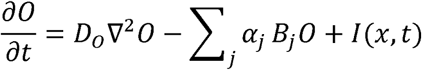

where:

- *D_o_* is oxygen diffusivity,
- *α_j_* represents species-specific oxygen utilization,
- *I*(*x,t*)represents environmental oxygen input.

The diffusion term generates spatial oxygen gradients, whereas biomass-dependent consumption continuously reshapes these gradients locally.

### 3.4. Radical field dynamics

The defining murburn assumption is that radical generation peaks under intermediate oxygen conditions rather than increasing monotonically with oxygen concentration.

We therefore define:

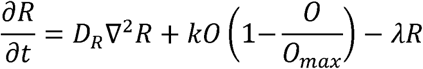

where:

- *D_R_* is effective radical diffusivity,
- *K* is radical-generation strength,
- *0_max_* represents the oxygen concentration above which radical quenching

dominates,

- *λ* represents radical decay and scavenging.

The nonlinear generation term:

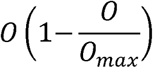

is maximal at intermediate oxygen tensions and minimal under both fully oxic and fully anoxic conditions. This equation mathematically encodes the central ecological proposition of the present work.

### 3.5. Biomass dynamics

Species biomass evolves according to:

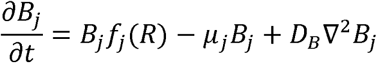

where:

- *f_j_*(*R*)is redox-dependent fitness,
- *µ_j_* is mortality,
- *D_B_* is dispersal diffusivity.

Fitness is assumed to depend non-monotonically upon radical intensity:

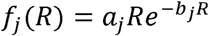

where:

- *a_j_* controls radical-utilization efficiency,
- *b_j_* controls oxidative sensitivity.

This structure reflects a biologically realistic trade-off:

- low radical intensity yields insufficient energetic activation,
- excessive radical exposure becomes damaging,
- intermediate radical flux maximizes viability.

The adopted fitness structure represents the simplest unimodal response capable of encoding both activation at low-to-intermediate DRS flux and inhibition under excessive oxidative exposure. Similar non-monotonic stress-response functions are widely employed in ecological and toxicological modeling. Different species possess distinct and values, thereby generating overlapping viability optima across redox space.

### 3.6. Emergence of coexistence

The murburn ecological formalism differs fundamentally from classical competition models because fitness is not globally fixed. Instead, local viability depends dynamically upon fluctuating redox conditions generated through coupled reaction–diffusion processes.

Several immediate consequences follow:

1. No single metabolic strategy remains universally optimal.
2. Local environmental structure continuously reshapes competition (Morris et al., 2012).
3. Organisms alter the same redox fields governing their fitness.
4. Spatial heterogeneity stabilizes coexistence.
5. Competitive exclusion becomes dynamically suppressed.

Thus, biodiversity emerges naturally from stochastic redox heterogeneity without requiring externally imposed coexistence rules or fine-tuned deterministic partitioning. Unlike classical Turing-type morphogen systems, the present framework does not require deterministic activator–inhibitor pairing or stationary pattern instability conditions. Instead, coexistence emerges from stochastic redox-dependent viability landscapes generated through diffusible oxidative intermediates and biologically coupled oxygen consumption.

### 3.7. Environmental engineering and ionic amplification

Aquatic ecosystems differ fundamentally from terrestrial systems because water-rich ionic environments spatially extend diffusional redox interactions through enhanced conductivity, halide-mediated chemistry, and constrained oxygen diffusion. Consequently, ecological organization in aquatic systems may depend not only upon local oxygen gradients but also upon organism-mediated restructuring of diffusional redox architectures. To incorporate such effects minimally, the oxygen equation may be extended by introducing an environmental-engineering term:

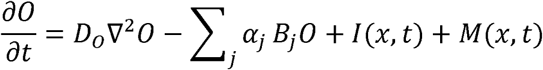

where:

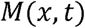

represents organism-mediated restructuring of local oxygenation and diffusional transport. Biologically, this term may represent: gill ventilation, mucus-mediated diffusion barriers, benthic sediment mixing, coral-associated interfacial restructuring, or biofilm-generated diffusional heterogeneity. Importantly, the environmental-engineering term does not represent direct trophic interaction. Instead, organisms indirectly influence coexistence by dynamically reshaping local redox landscapes.

Aquatic ionic environments may further amplify diffusional radical coupling. This effect may be incorporated phenomenologically by allowing effective radical diffusivity to depend upon ionic conductivity:

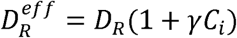

where:

- *C_i_* represents effective ionic strength,
- and *γ* represents conductivity-dependent enhancement of diffusional redox coupling.

This term reflects the possibility that halide-rich aqueous systems spatially extend reactive-field propagation through radical relay chemistry, ionic stabilization, and interfacial electrochemical interactions.

Under these conditions, aquatic ecosystems become capable of generating ESDRA in which localized radical chemistry acquires spatial ecological significance far beyond immediate reaction sites. Importantly, these additions preserve the minimal structure of the murburn ecological formalism while extending its applicability to aquatic systems characterized by strong ionic coupling and organism-mediated environmental restructuring.

## 4. Simulations

### 4.1. Objectives and rationale

The simulations developed herein aim to determine whether a minimal murburn-based reaction–diffusion framework naturally produces:

1. radical maxima at intermediate oxygen tensions,
2. stable coexistence among competing metabolic strategies,
3. biodiversity peaks within micro-oxic regions,
4. and dynamically shifting ecological mosaics.

Importantly, the simulations are intentionally minimal. The objective is not to reproduce every biochemical detail of real ecosystems, but to determine whether the central murburn proposition alone is sufficient to generate experimentally observed ecological behaviors.

The central proposition tested is: *Intermediate oxygen tensions maximize stochastic redox heterogeneity and DRS flux, thereby generating dynamically shifting fitness landscapes that suppress competitive exclusion and promote coexistence*.

Reaction–diffusion systems are particularly appropriate for such investigations because ecological microenvironments are fundamentally diffusion-limited and spatially heterogeneous (Levin, 1992; Murray, 2002). Oxygen gradients in sediments, gut mucus layers, rhizospheres, and OMZ interfaces emerge from the interplay between diffusion and localized biological consumption (Albenberg et al., 2014; Wright et al., 2012). Simultaneously, extracellular ROS production and radical-mediated chemistry introduce nonlinear feedback between environmental chemistry and organismal viability (Diaz et al., 2013; Hansel et al., 2019). The simulations therefore examine whether: oxygen diffusion, nonlinear radical generation, and redox-dependent fitness are alone sufficient to generate stable biodiversity maxima within micro-oxic systems.

### 4.2. One-dimensional oxygen-gradient simulations

The simplest physically meaningful configuration consists of a one-dimensional spatial domain extending from an oxygen-rich boundary toward an oxygen-poor region. This configuration approximates: gut mucus gradients, sediment interfaces, microbial mats, and oxygen-penetration fronts in soils.

Oxygen is introduced from one boundary and allowed to diffuse inward while simultaneously being consumed by biological populations. Radical generation is coupled nonlinearly to local oxygen concentration according to the murburn formalism developed in Section 3, as shown in Figure 1 (MATLAB Code for the same is given in Supplementary Information, Item 1). Under these conditions: oxygen declines spatially, reactive intermediates peak away from both oxic and anoxic extremes, and different species stabilize preferentially at distinct radical intensities.

**Figure 1:**
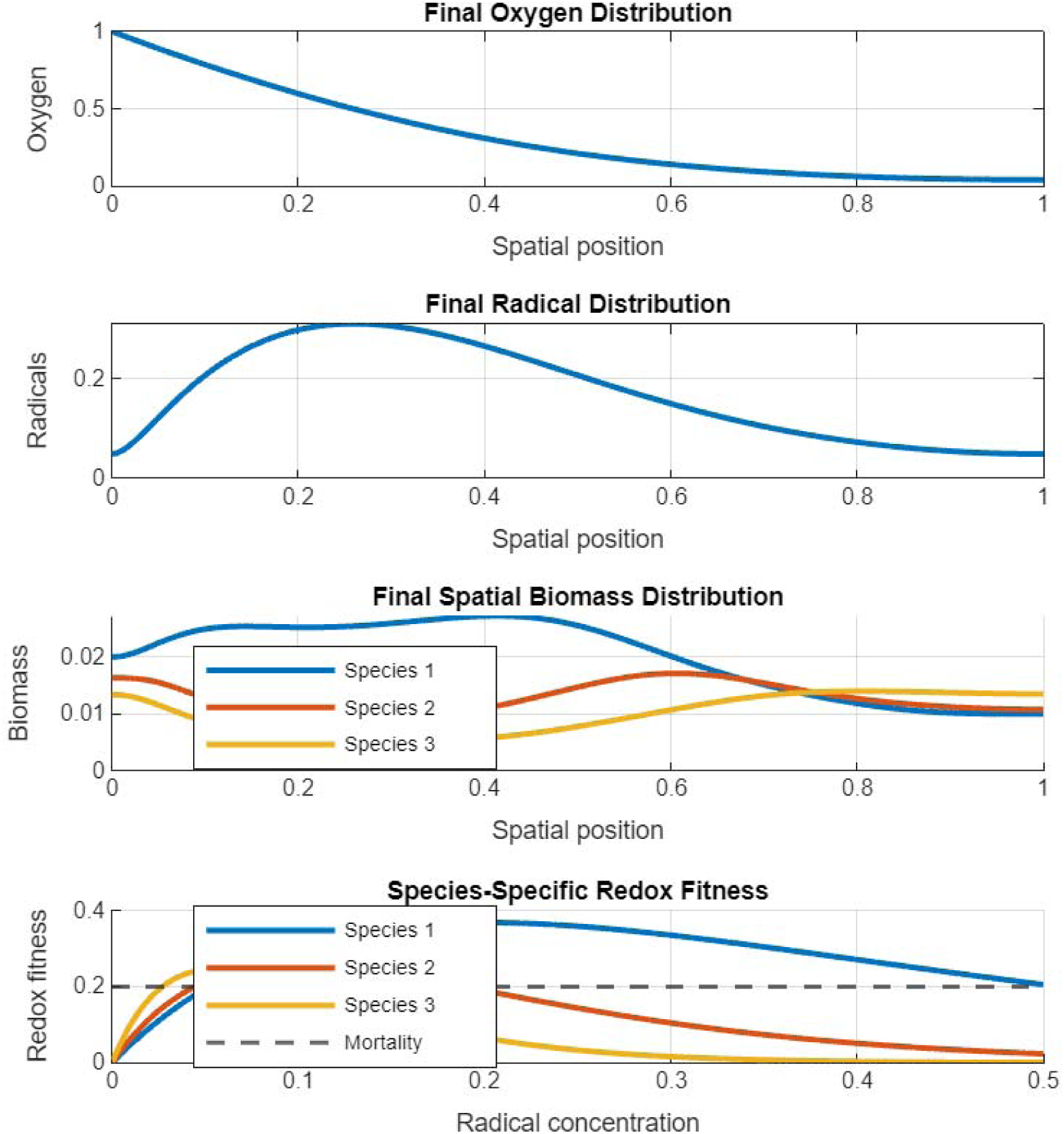
1D simulation of biomass variability as a function of spatial variability of oxygen and DRS.

Species possessing lower oxidative tolerance initially dominate near highly oxic regions but decline as radical quenching suppresses productive redox heterogeneity. Conversely, organisms requiring higher radical activation fail within strongly anoxic regions because insufficient redox throughput exists to sustain activation. Intermediate radical specialists therefore stabilize within micro-oxic domains. Importantly, biodiversity maxima emerge spontaneously without: externally imposed niche partitioning, deterministic coexistence rules, or finely tuned interaction matrices. This result differs fundamentally from classical Lotka–Volterra systems, which generally require carefully balanced interaction coefficients to maintain coexistence (Lotka, 1925; Volterra, 1926). A comparison with the Lotka-Volterra model is shown (along with the MATLAB Code) in Supplementary Information, Item 2.

### 4.3. Macroscopic-fauna-mediated redox landscape engineering (1D)

An important implication of the murburn ecological framework is that organisms may indirectly promote biodiversity by restructuring local redox architectures rather than solely through direct trophic or competitive interactions.

Aquatic macrofauna continuously modify: oxygen penetration, mucus-layer diffusion, ionic microenvironments, particle transport, and extracellular oxidative chemistry. Examples include: gill ventilation, benthic burrowing, coral mucus secretion, filter feeding, and biofilm restructuring. Under murburn assumptions, such processes alter local oxygen gradients and diffusional radical propagation, thereby reshaping the ecological viability landscape itself. To investigate this possibility, the reaction–diffusion framework was extended to include a moving macrofaunal environmental-engineering term. The oxygen equation becomes:

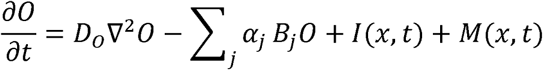

where:

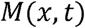

represents macrofauna-mediated oxygenation and diffusional restructuring.

This term may represent: localized oxygen injection, mucus-mediated diffusion barriers, sediment mixing, or interfacial ionic modification. Importantly, the macrofaunal organisms are not modeled as dominant competitors. Instead, they function as environmental redox architects whose activities dynamically reshape ecological conditions for surrounding microbial and mesoscopic communities. Simulations demonstrate that moving oxygenation interfaces generate persistent radical hotspots and dynamically shifting coexistence zones (Figure 2a and 2b; MATLAB Code for the same is given in Supplementary Information, Item 3). Biodiversity becomes concentrated near engineered micro-oxic boundaries rather than uniformly distributed throughout the environment. This behavior differs fundamentally from classical ecological frameworks because environmental structure is no longer externally imposed. Instead, ecological architecture emerges through reciprocal coupling between organisms and diffusional redox fields.

**Figure 2 a and 2b:**
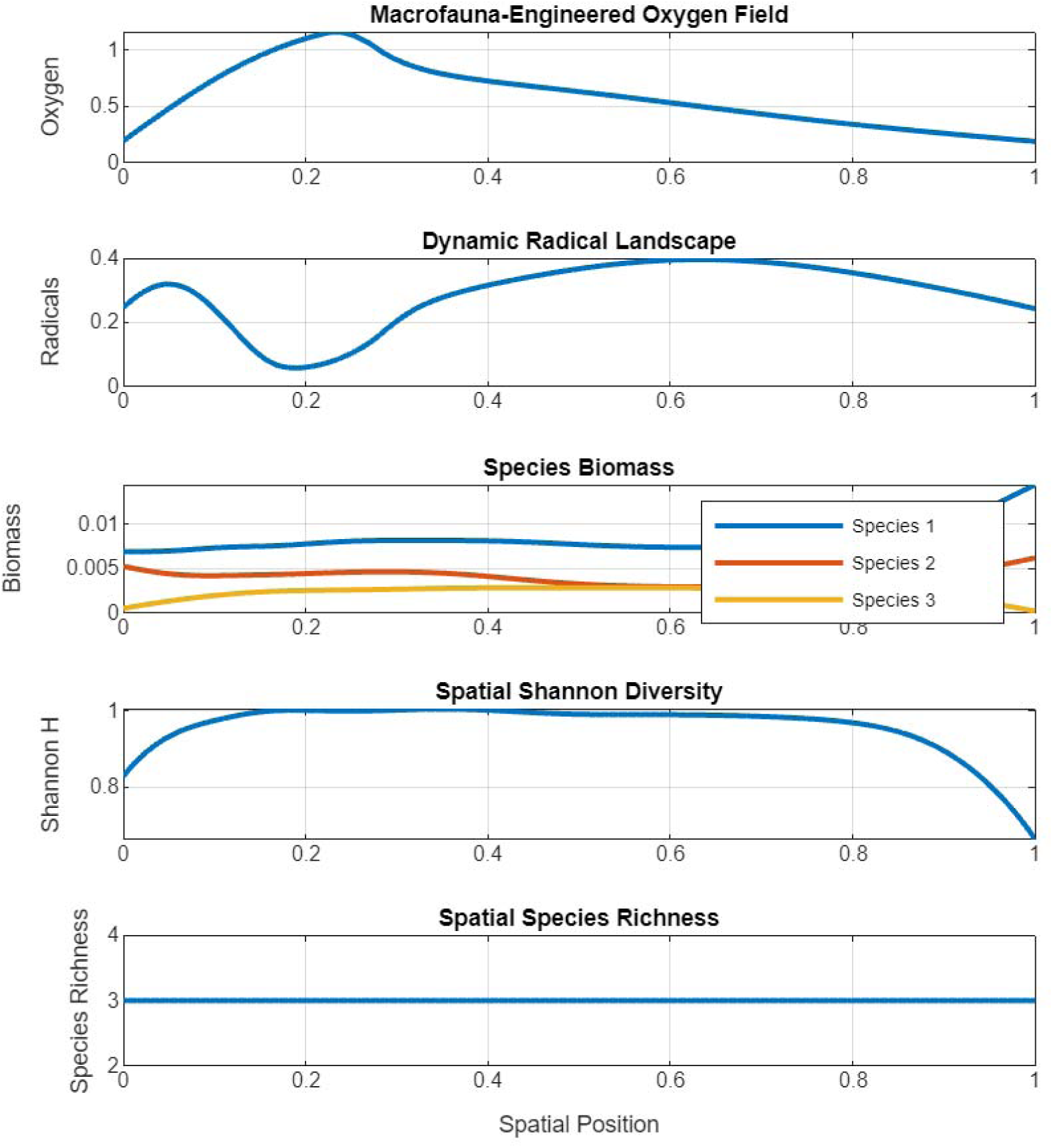
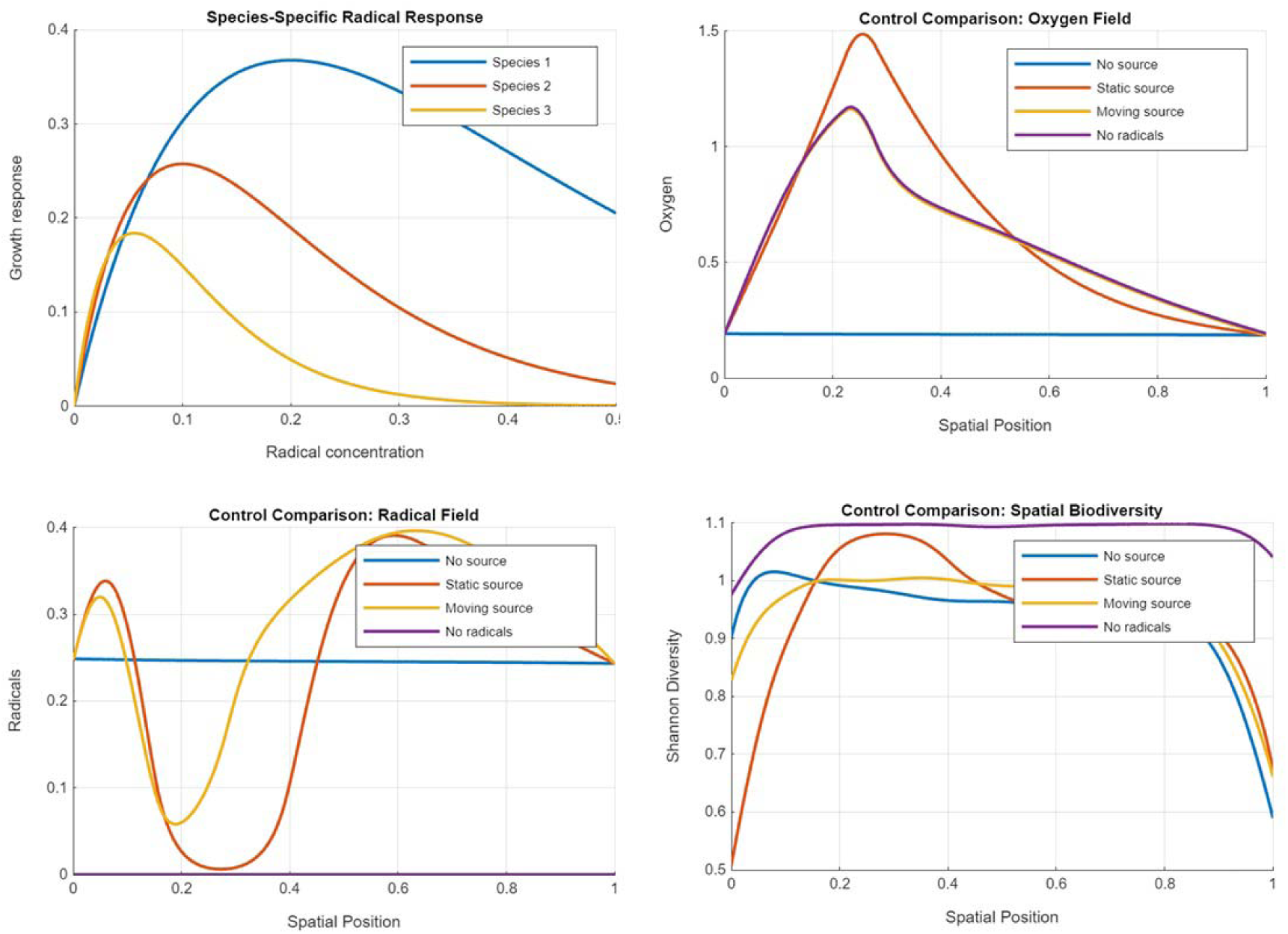
1D simulation of macrofauna-induced oxygen variations.

We disclaim clearly that the simulations above are qualitative exploratory demonstrations rather than quantitatively parameterized environmental reconstructions. Parameter choices were selected to maintain numerical stability and illustrate emergent coexistence behavior across broad regions of parameter space.

### 4.4. Two-dimensional redox mosaics

Real ecosystems are not one-dimensional systems but spatially heterogeneous redox mosaics. Rhizospheres, sediment interfaces, gut mucus layers, coral microbiomes, and marine particle aggregates all contain irregular oxygen distributions shaped by: diffusion, hydration, local metabolism, extracellular polymer networks, and fluid transport. To approximate such environments, the murburn framework was extended into two spatial dimensions. The MATLAB Code used for the same (Supplementary Information, Item 4) incorporates: multiple macroflora oxygen sources, multiple macrofauna oxygen hotspots, fractal-like environmental heterogeneity, murburn DRS generation, five species with differing DRS preferences, dominant species map and Shannon diversity map. Under these conditions: multiple oxygen sources generates irregular gradients, radical hotspots emerge dynamically, biomass distributions self-organize spatially, and coexistence islands form spontaneously. Importantly, biodiversity no longer appears as smooth stratification alone but as dynamically shifting ecological patches. These simulations (Figure 3) reproduce several experimentally observed ecological properties: patch formation, transient coexistence fronts, fluctuating dominance, and localized diversification zones. Such behaviors resemble microbial patterning observed in: biofilms, rhizospheric networks, OMZ interfaces, and microbial mats (Levin, 1992; Falkowski et al., 2008).

**Figure 3:**
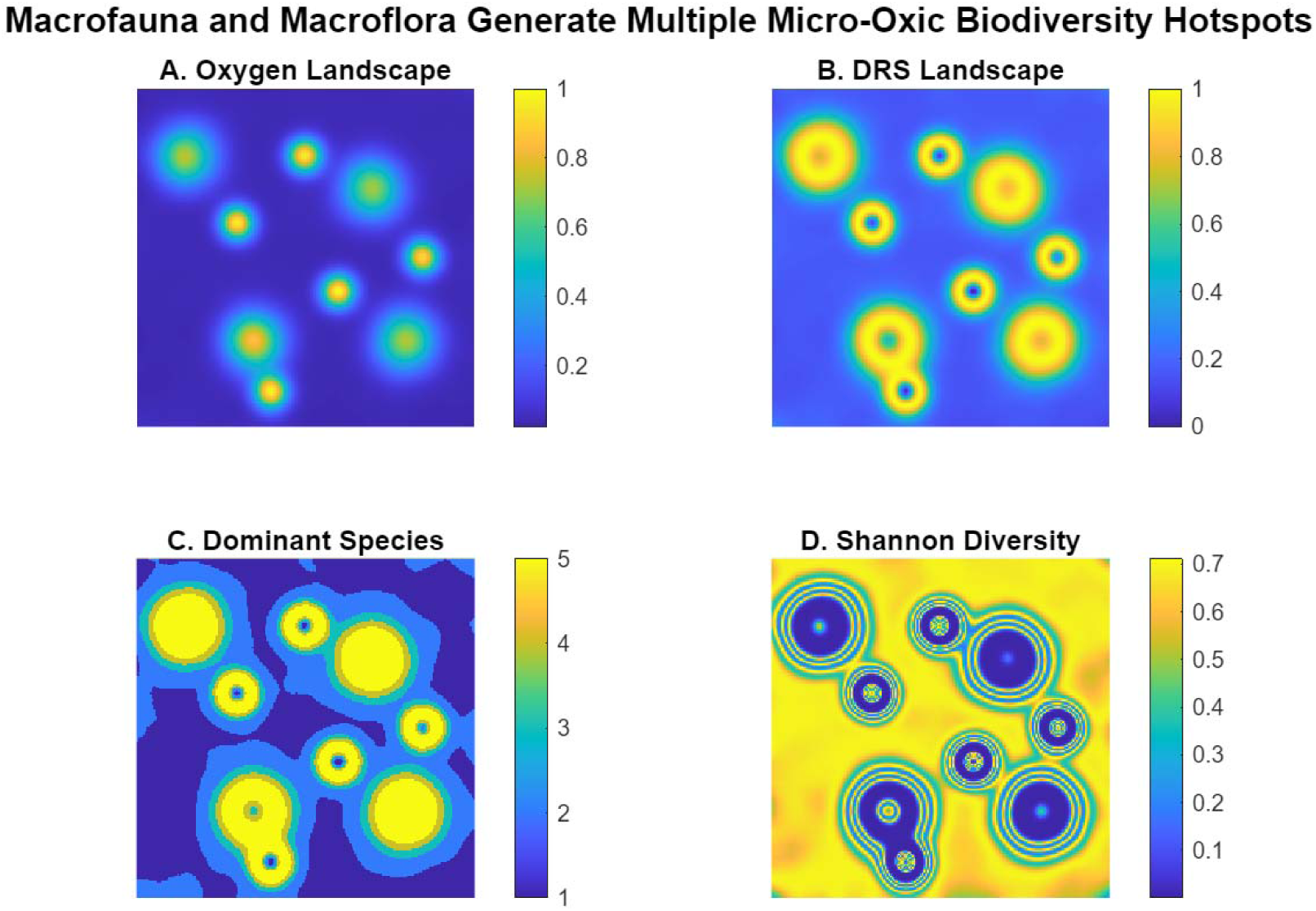
Panel A (Oxygen Landscape): irregular oxygen plumes generated by macroflora and macrofauna, modulated by environmental heterogeneity. Panel B (DRS Landscape): patchy micro-oxic halos and interconnected redox corridors where DRS is highest. Panel C (Dominant Species): ecological niche partitioning according to preferred DRS regime. Panel D (Shannon Diversity): multiple biodiversity hotspots and corridors, illustrating that diversity peaks at intermediate oxygen / high redox heterogeneity rather than at oxygen maxima.

### 4.4. Evolutionary extension

An important consequence of murburn ecology is that mutation and diversification need not occur uniformly across space. Reactive intermediates are known to influence: DNA damage, oxidative modification, transposition, epigenetic alteration, and stress-associated mutagenesis (Manoj, 2025). Accordingly, the model permits species-specific oxidative sensitivity parameters to evolve according to local radical intensity:

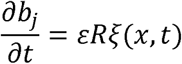

where:

- ɛ defines evolutionary timescale,
- and ξ(*x,t*)represents stochastic mutational direction.

Under these conditions: adaptation occurs fastest near intermediate radical zones, diversification localizes within fluctuating redox interfaces, and evolutionary innovation concentrates within micro-oxic regions. Thus, micro-oxic environments behave simultaneously as: ecological coexistence zones, mutational hotspots, and potential zones of elevated adaptive turnover.

## 5. Predictions, falsifiability and empirical corroboration

The murburn ecological framework predicts that biodiversity should correlate more strongly with redox heterogeneity, oxygen fluctuation, and radical-field variability than with mean oxygen concentration alone. Several independent ecological systems support this prediction. Marine oxygen minimum zones exhibit maximal microbial functional diversity near fluctuating oxic–anoxic boundaries rather than within uniformly oxic waters (Wright et al., 2012). Extracellular ROS production by marine microbes correlates strongly with zones of intense ecological turnover and nutrient cycling (Diaz et al., 2013; Hansel et al., 2019). In gut ecosystems, oxygen gradients strongly determine microbiome partitioning, with moderate oxidative interfaces supporting coexistence, whereas severe inflammation or complete anoxia both reduce microbial complexity (Albenberg et al., 2014; Zheng et al., 2015). Rhizospheric and soil aggregate systems likewise exhibit maximal diversity in fluctuating hydration-dependent micro-oxic domains where metal-mediated radical chemistry remains active (Kuzyakov & Blagodatskaya, 2015)

Importantly, the framework proposes that fluctuating radical fields themselves constitute ecological structuring mechanisms (continuously reshaping fitness landscapes, suppressing competitive exclusion, and enabling stable coexistence) rather than merely stating that “ROS matter biologically.” Increasing evidence indicates that aquatic macrofauna actively restructure environmental oxygen gradients and associated redox chemistry. Bioturbation alters sediment oxygen penetration, coral mucus generates chemically active interfacial layers (Wild et al., 2004; Glasl et al., 2016), and activities such as gill ventilation, filter feeding, burrowing, and mucus secretion continuously reshape local oxygen distributions and diffusional transport. Marine snow particles and biofilms further demonstrate how biologically generated diffusional barriers create localized oxygen-depletion zones surrounded by oxygen-rich waters, supporting extraordinary microbial diversity despite spatial confinement (Azam & Malfatti, 2007). Under the murburn framework, these observations are interpreted as organism-mediated redox landscape engineering; extending niche construction theory into stochastic redox ecology (Laland et al., 2016; Laland et al., 2017).

### 5.1. Biodiversity should correlate with oxygen variance rather than mean oxygen concentration

Classical models predict monotonic responses to oxygen; murburn ecology predicts that uniformly oxic and uniformly anoxic systems both support lower biodiversity than fluctuating micro-oxic interfaces. This is experimentally testable via microcosms manipulating oxygen variance independently of mean concentration.

### 5.2. Intermediate ROS buffering should maximize coexistence

Insufficient buffering leads to oxidative collapse; excessive buffering suppresses productive redox heterogeneity. Moderate buffering should maximize species richness: testable in microbial consortia, gut microbiomes, and biofilms by manipulating antioxidant capacity.

### 5.3. Artificial homogenization of oxygen gradients should reduce biodiversity

Forced homogenization (e.g., via mixing, aeration, or microfluidic control) should suppress coexistence and drive competitive exclusion. This prediction is directly testable in sediments, gut models, and rhizosphere systems.

### 5.4. Evolutionary diversification should peak in fluctuating redox interfaces

Since reactive intermediates influence mutagenesis and stress-associated variation, the model predicts elevated genomic variability, adaptive turnover, and innovation within OMZ boundaries, rhizospheres, gut mucus layers, and microbial mats.

### 5.5. Halide-rich marine systems should exhibit stronger diffusional redox structuring than freshwater systems

Seawater’s high ionic strength and halide abundance (Cl□, Br□, I□) promote halogen-radical relay chemistry and interfacial oxidative coupling. Thus, marine ecosystems should show more persistent and spatially extended radical-field coupling than freshwater systems, which should exhibit shorter-range coupling and greater sensitivity to oxygen collapse: testable via comparative oxygen microprofiling and ROS mapping across marine and freshwater interfaces.

Collectively, these predictions are falsifiable: if biodiversity correlated monotonically with oxygen, if homogenization failed to reduce diversity, or if marine–freshwater redox structuring proved equivalent, the murburn framework would require substantial revision.

## 7. Conclusion

The present work proposes that biodiversity is fundamentally coupled to the spatial and temporal organization of redox heterogeneity. Rather than treating oxygen solely as a monotonic energetic resource, the murburn ecological framework interprets fluctuating oxygen fields as generators of diffusible reactive intermediates that dynamically restructure ecological viability landscapes. Under such conditions, coexistence emerges not from static deterministic partitioning alone, but from continuously shifting radical-mediated microenvironments that prevent stable global dominance by any single metabolic strategy.

The central result of the present study is that intermediate oxygen tensions naturally maximize stochastic redox heterogeneity. This generates localized radical fluxes capable of simultaneously supporting metabolic activation, imposing oxidative constraints, reshaping signaling dynamics, and influencing adaptive turnover. Consequently, fluctuating micro-oxic systems become chemically self-organizing coexistence zones in which ecological interactions emerge from coupled reaction–diffusion dynamics rather than fixed interaction matrices. The salient message of the current writing is captured in Figure 4 below:

**Figure 4.**
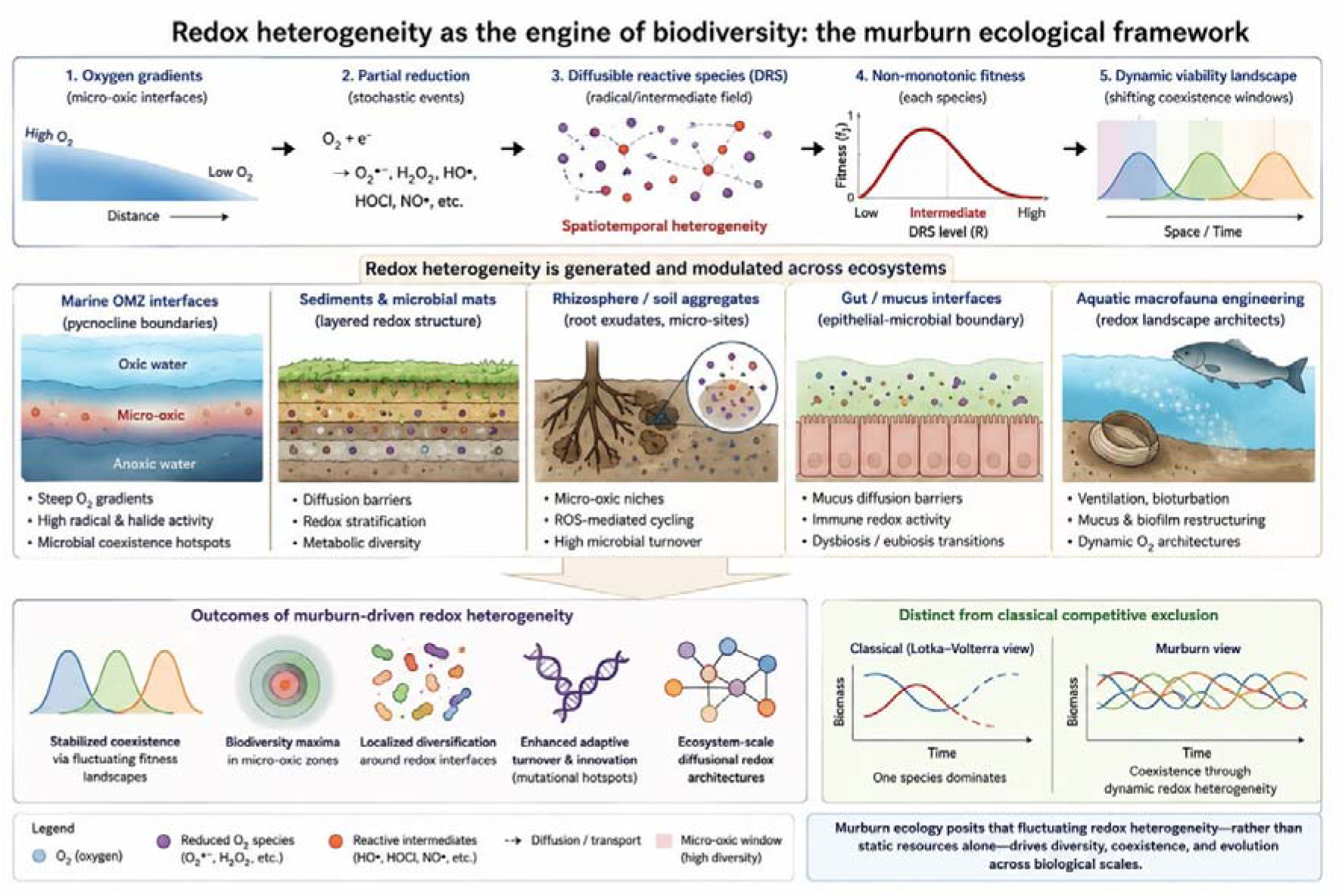
Conceptual schematic of murburn-driven biodiversity in micro-oxic ecosystems. Fluctuating oxygen gradients within aquatic and sedimentary environments generate spatially heterogeneous DRS (redox intermediates) fields through partial oxygen reduction and coupled redox cycling. Intermediate oxygen tensions maximize stochastic redox heterogeneity, producing dynamic coexistence zones that suppress stable competitive exclusion and promote biodiversity. Water-rich ionic environments, particularly marine systems enriched in halides and transition metals, spatially extend redox interactions through diffusional and interfacial coupling. Macroscopic aquatic organisms further act as redox landscape engineers by reshaping oxygen penetration, mucus-associated diffusion barriers, and local oxidative microenvironments. Together, these processes generate ESDRA linking molecular redox dynamics to ecological coexistence, adaptive turnover, and biodiversity maintenance across micro-oxic interfaces.

Importantly, the framework further suggests that aquatic ecosystems amplify the ecological consequences of murburn-compatible dynamics because water-rich ionic environments spatially extend diffusional redox interactions. Marine systems, in particular, possess physicochemical properties (including persistent oxygen gradients, halide-rich chemistry, ionic conductivity, and interfacial transport limitations) that favor formation of ecosystem-scale redox architectures. Within such environments, macroscopic organisms may function as redox landscape engineers whose activities continuously restructure oxygen penetration, diffusional barriers, and radical-field heterogeneity.

The implications extend beyond theoretical ecology. Marine oxygen minimum zones, coral-associated interfaces, rhizospheric aggregates, sediments, microbial mats, biofilms, and gut mucus systems may all be interpreted as distributed stochastic redox reactors in which ecological organization emerges from chemically mediated heterogeneity. More broadly, the murburn ecological formalism provides a mechanistically integrated framework linking molecular redox dynamics, environmental chemistry, ecological coexistence, and evolutionary diversification across biological scales.

### Limitations

The present formulation intentionally isolates redox heterogeneity as a primary ecological structuring variable. Nutrient limitation, trophic coupling, and substrate-specific metabolic constraints are incorporated only implicitly. Future extensions may integrate explicit carbon, nitrogen, sulfur, and phosphorus fields into the reaction–diffusion framework.

## Supplementary Information

### Item 1: 4.2.1. MATLAB implementation: 1-D murburn biodiversity model

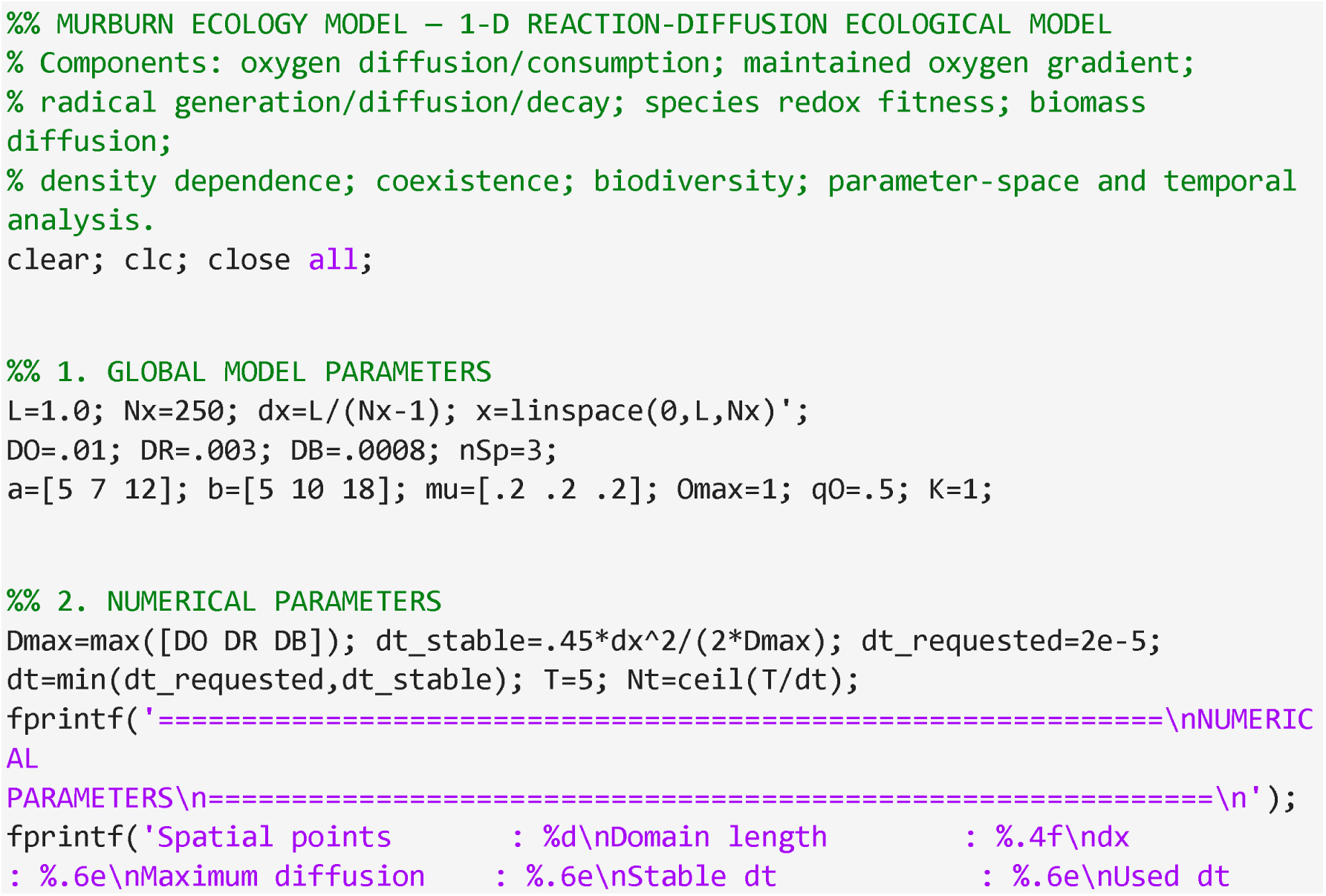
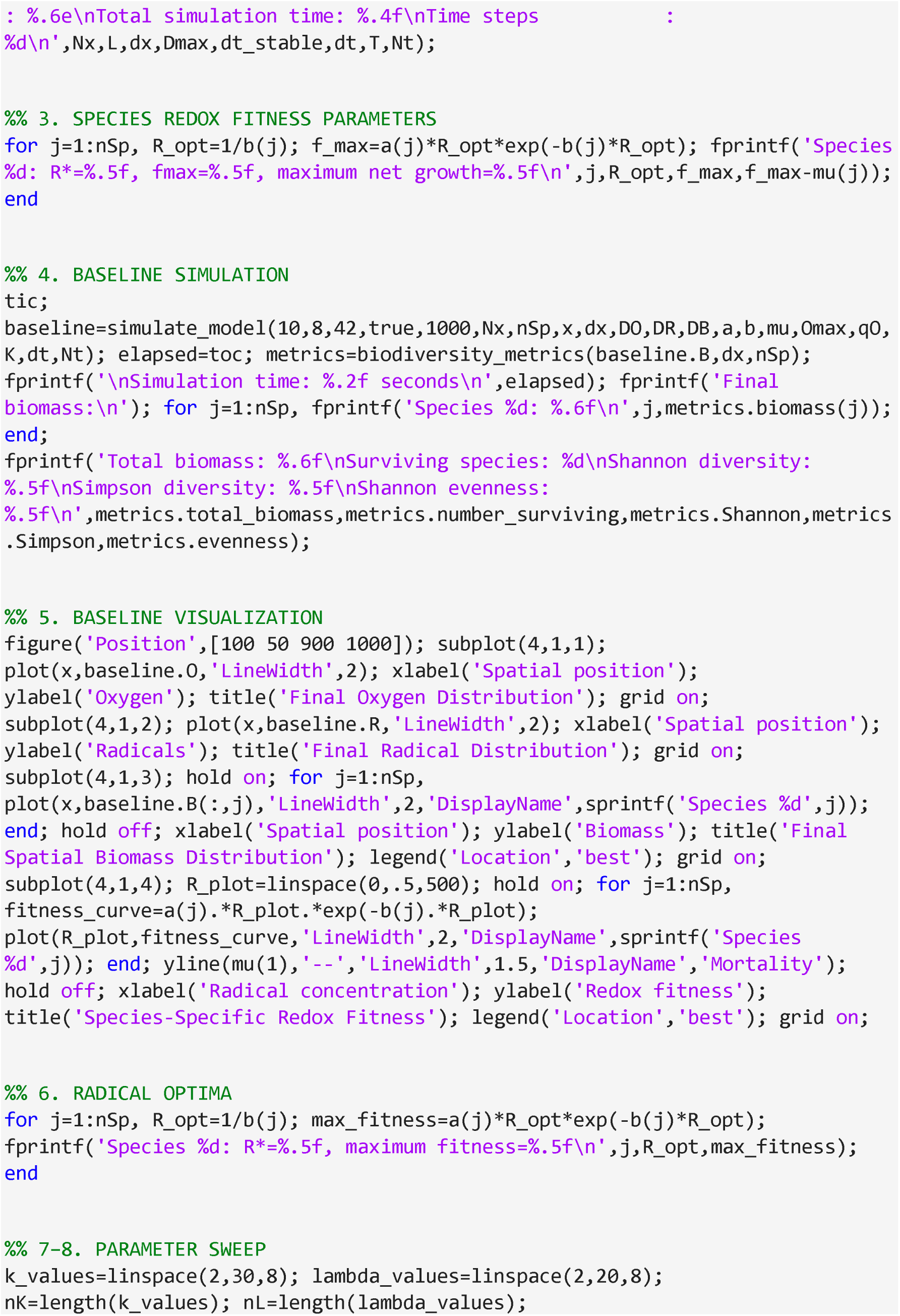
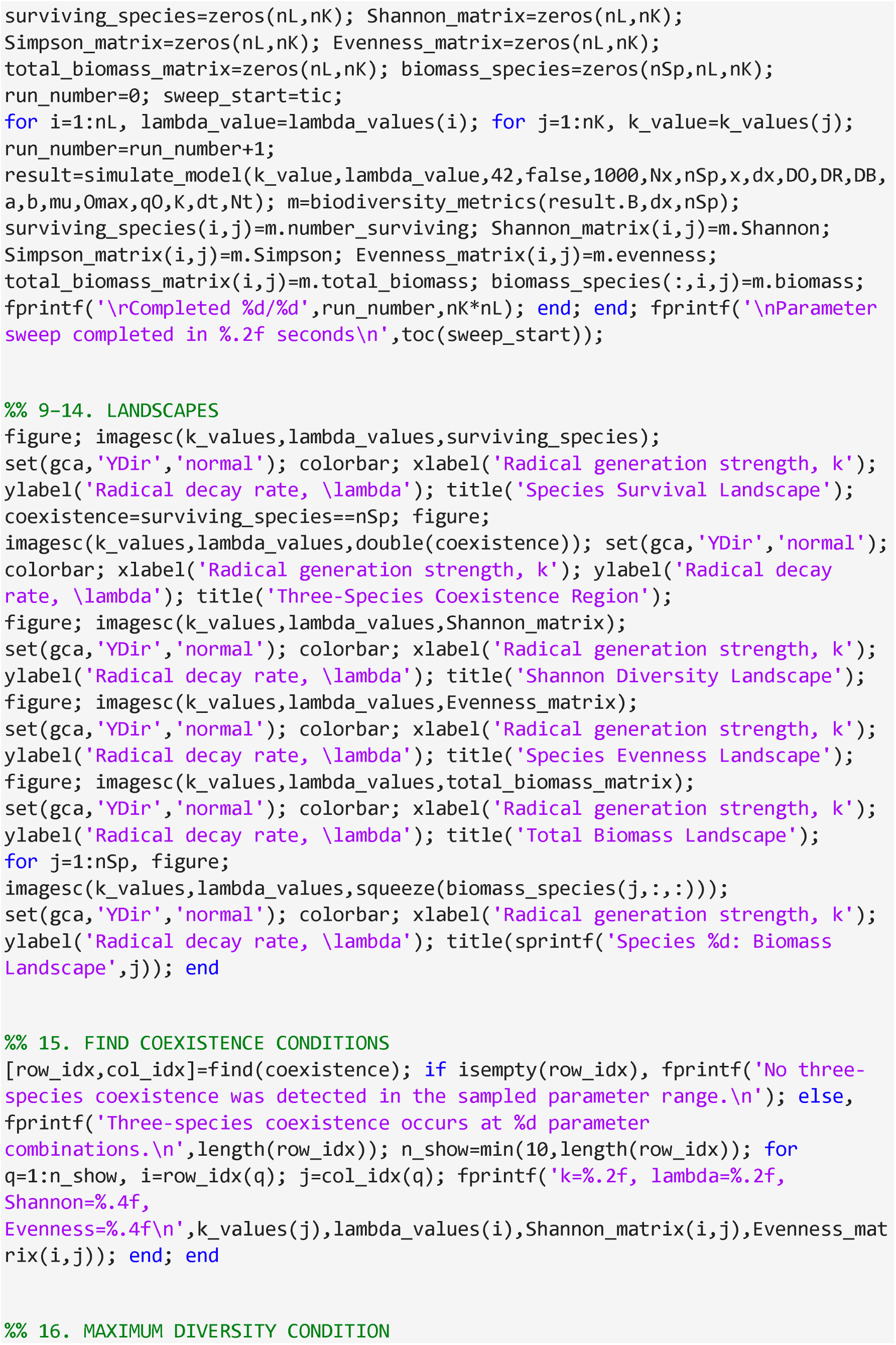
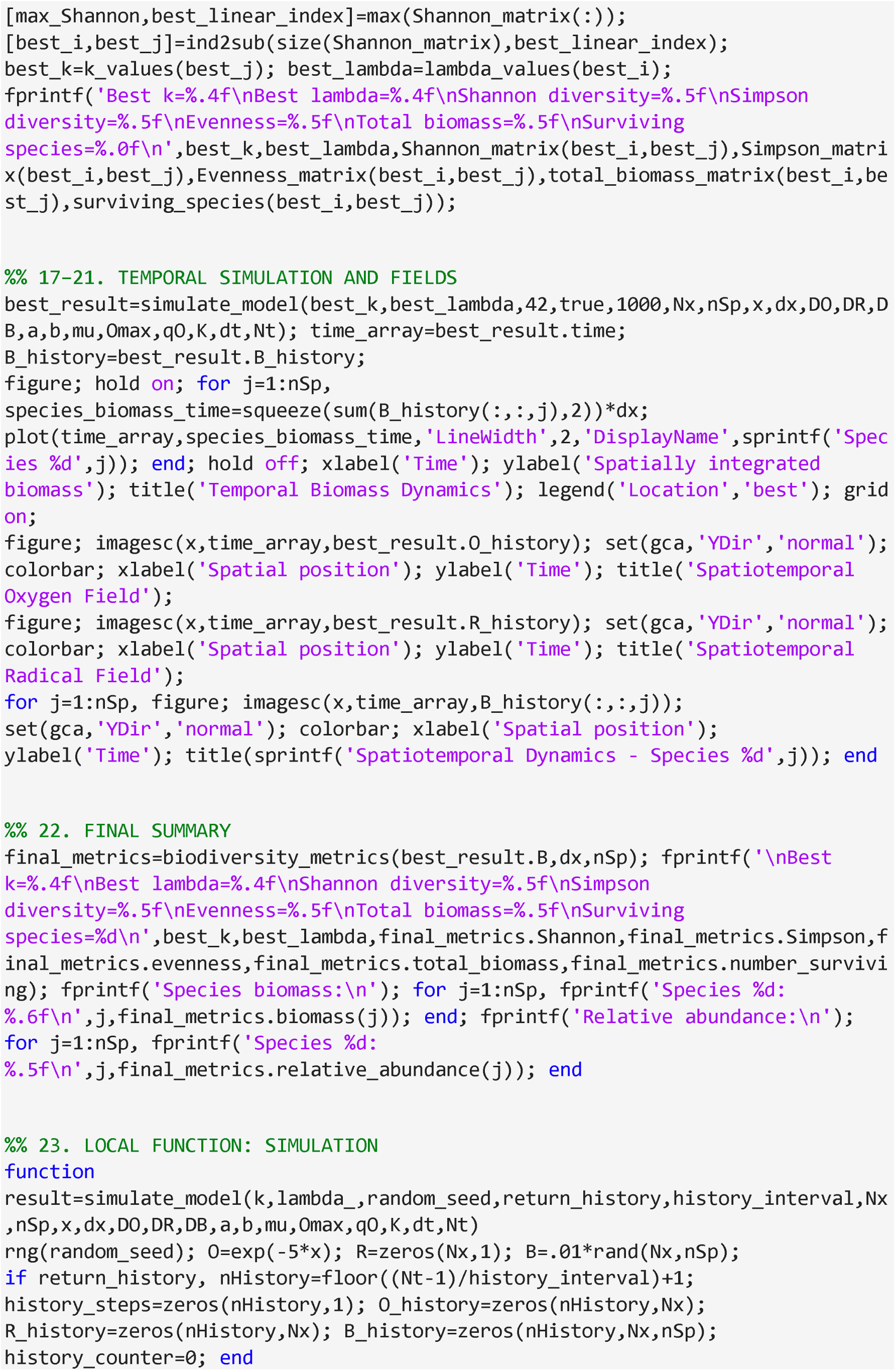
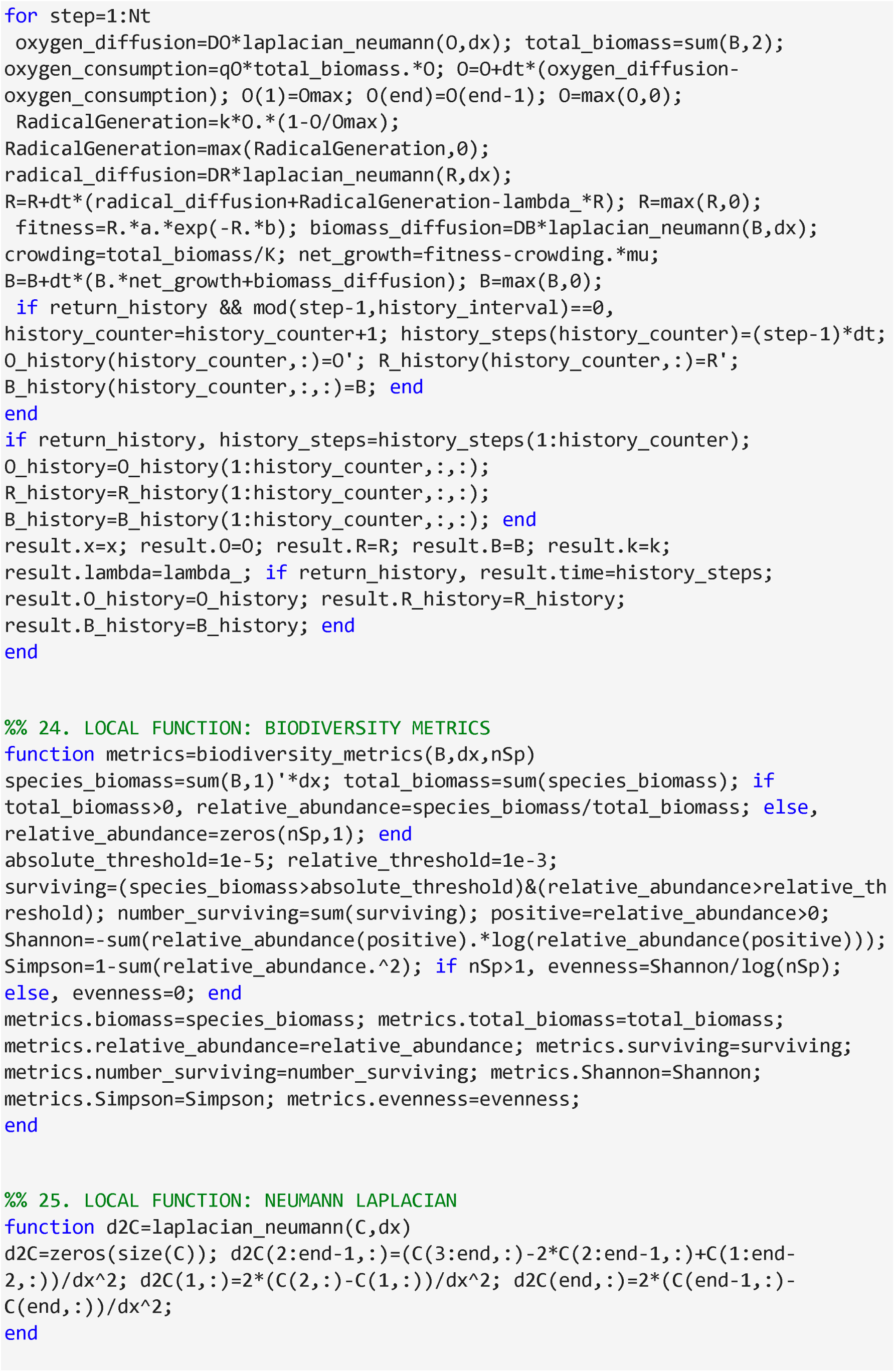

### Item 2: 4.2.2 Comparison with classical Lotka–Volterra dynamics

Classical Lotka–Volterra systems assume:

- fixed interaction coefficients,
- well-mixed populations,
- and static fitness ordering (Lotka, 1925; Volterra, 1926).

Under such conditions, competitive exclusion dominates unless coexistence parameters are carefully balanced.

The murburn ecological framework differs fundamentally because:

- fitness depends dynamically upon radical intensity,
- radical intensity depends upon oxygen gradients,
- and organisms reshape their own viability landscapes. Thus interaction coefficients become emergent rather than fixed.

#### MATLAB comparison: Lotka–Volterra competitive exclusion

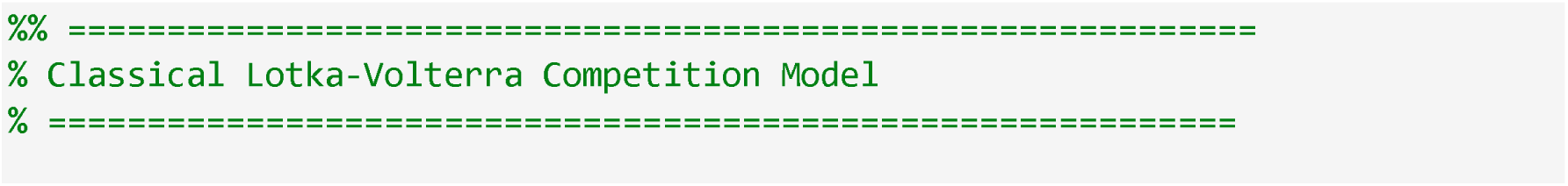
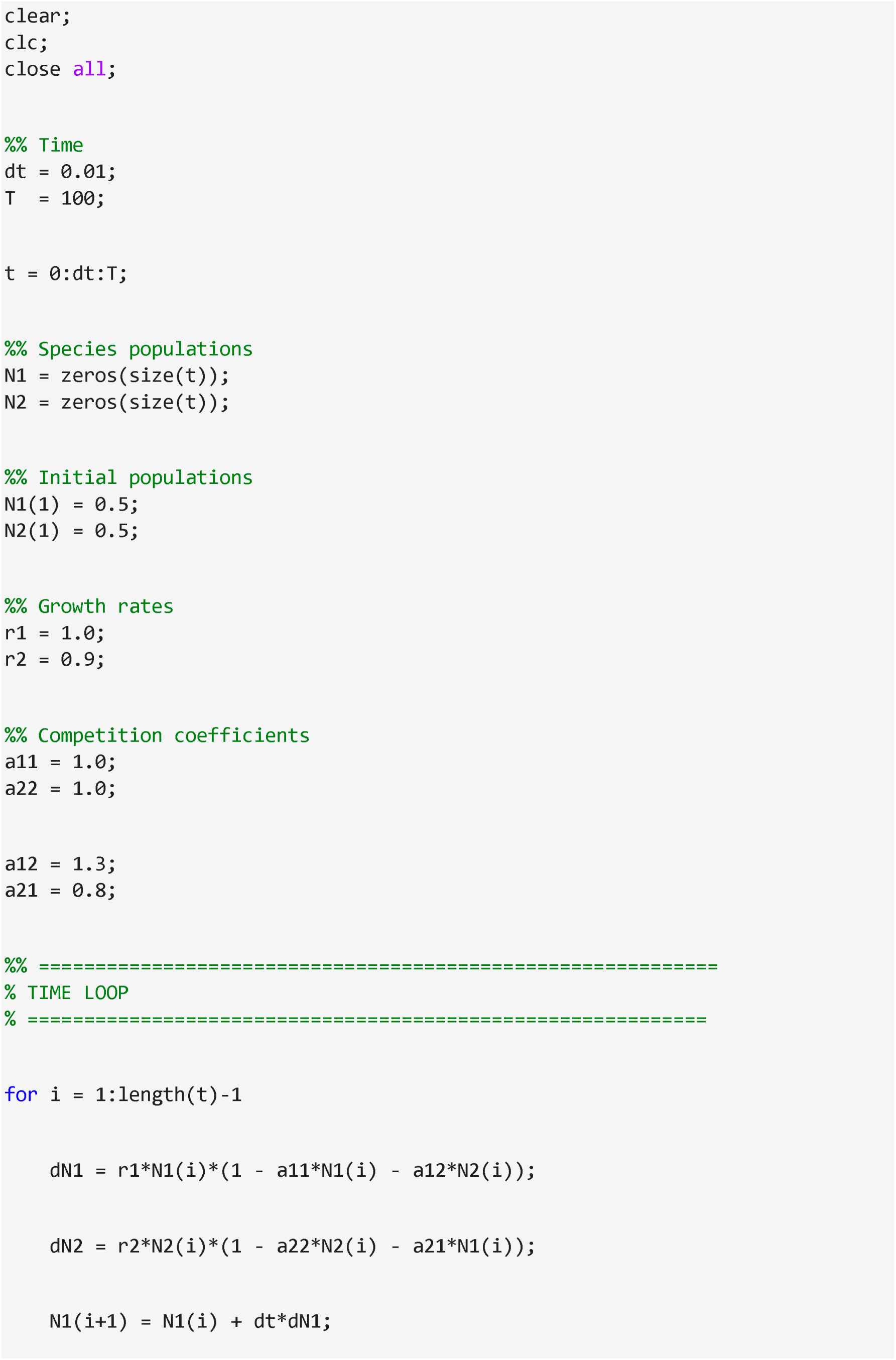
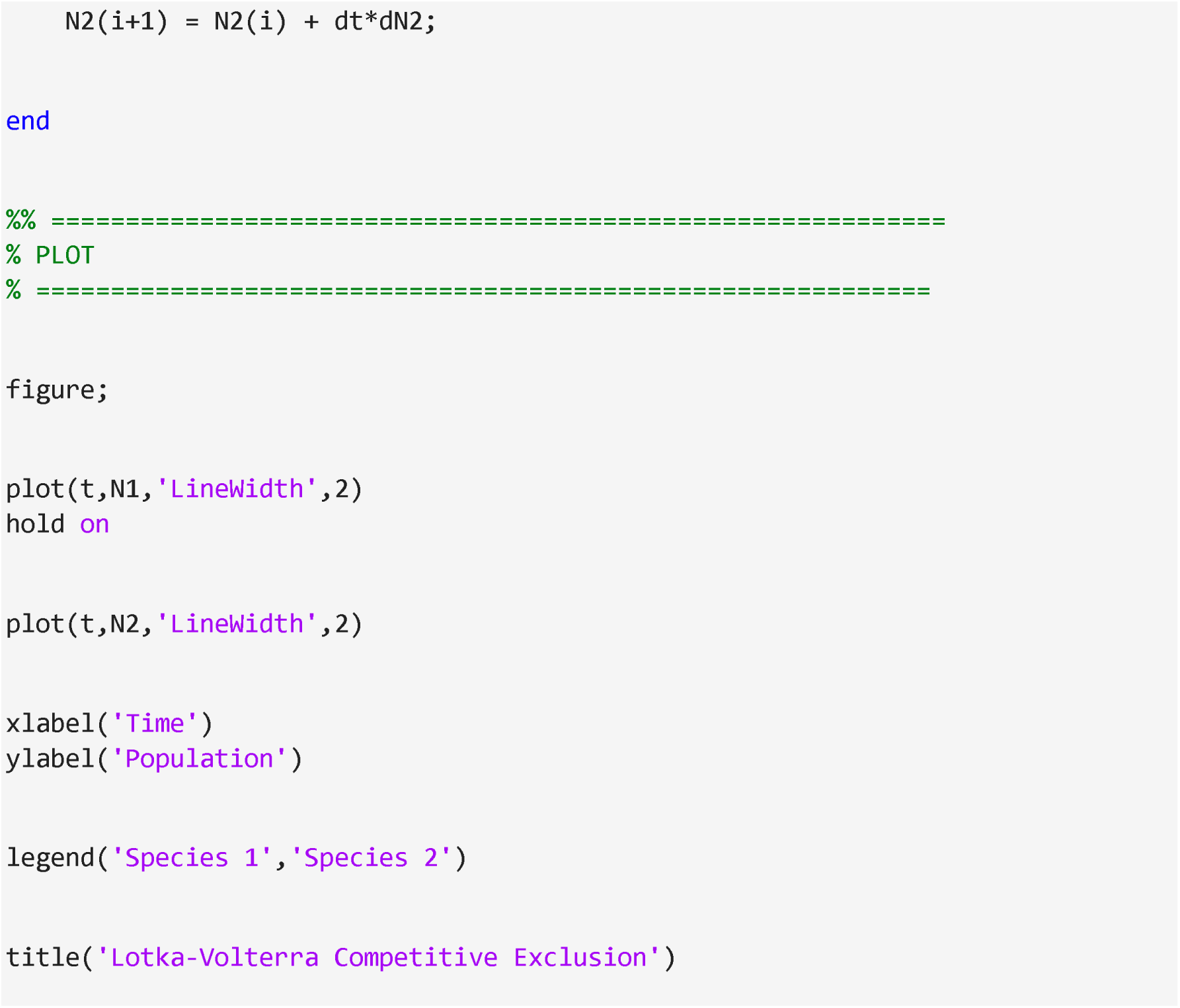

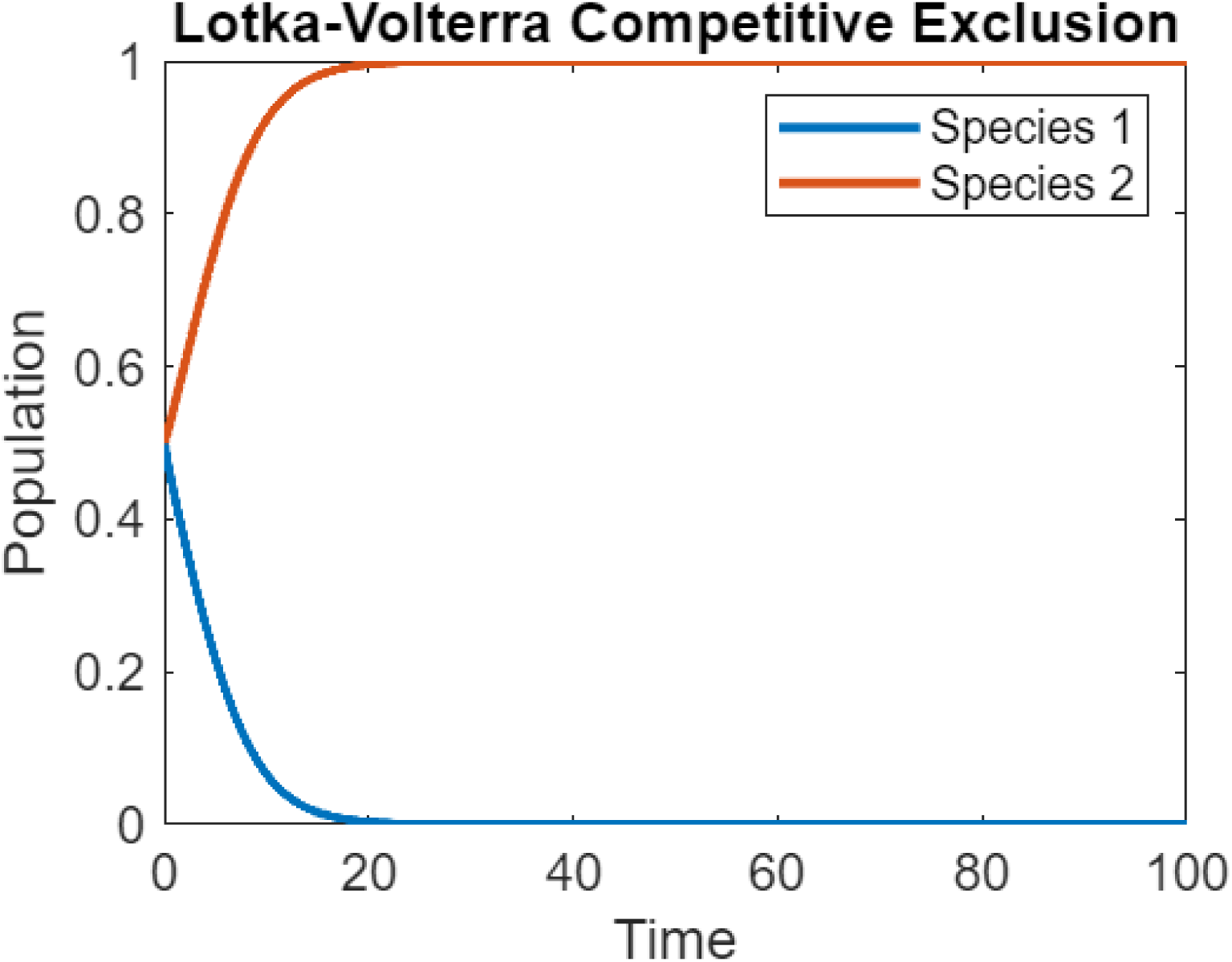

### Item 3: 4.3.1 MATLAB code for macroscopic fauna induced redox alterations

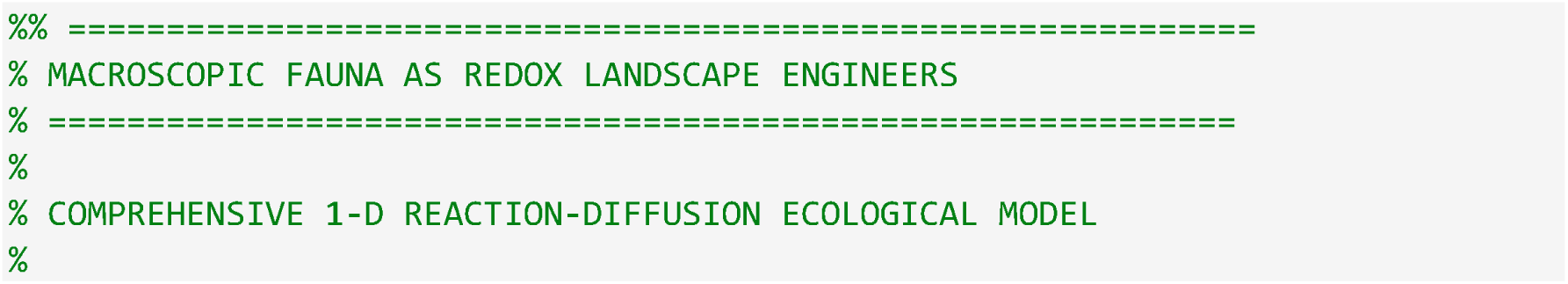
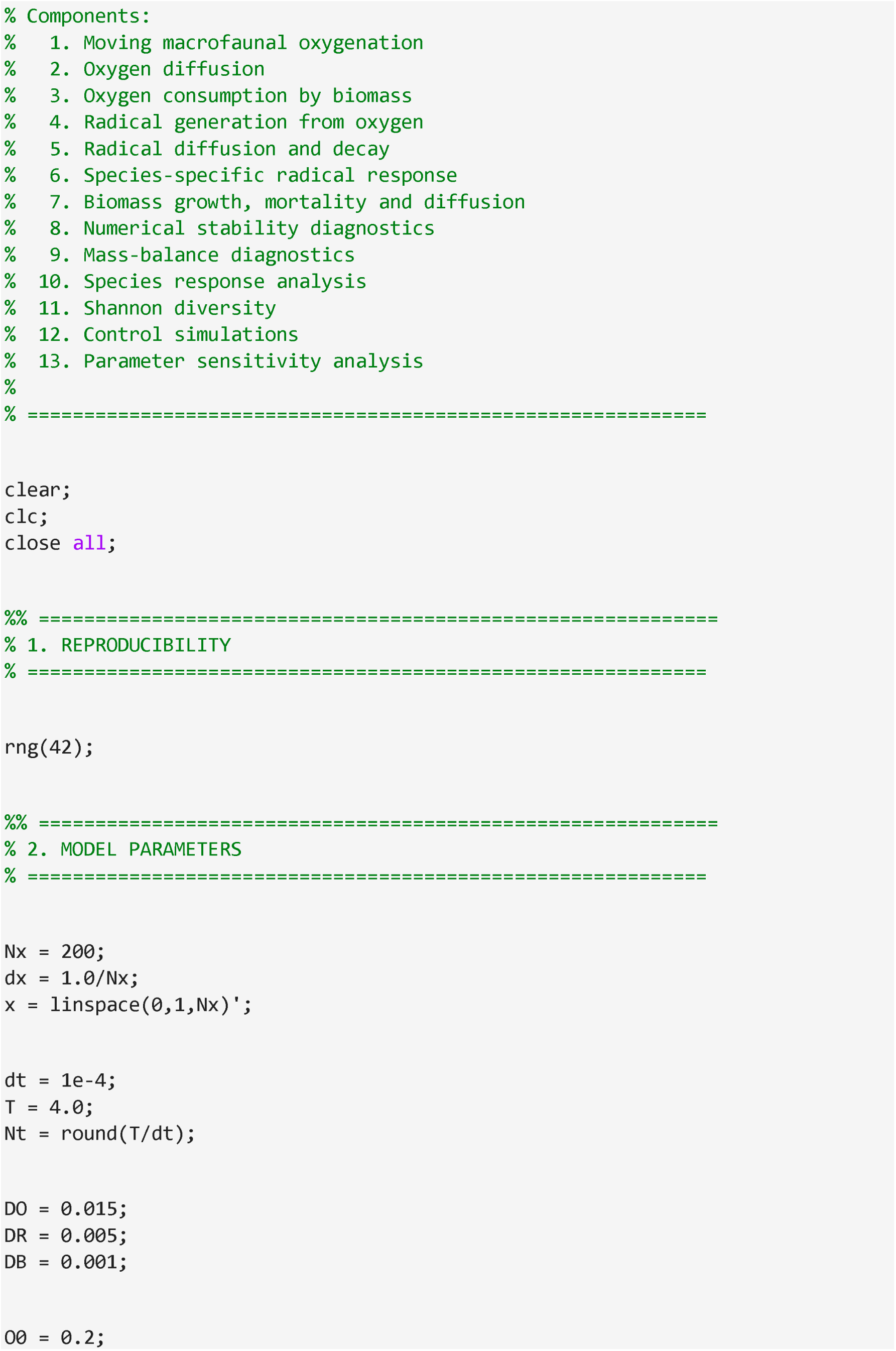
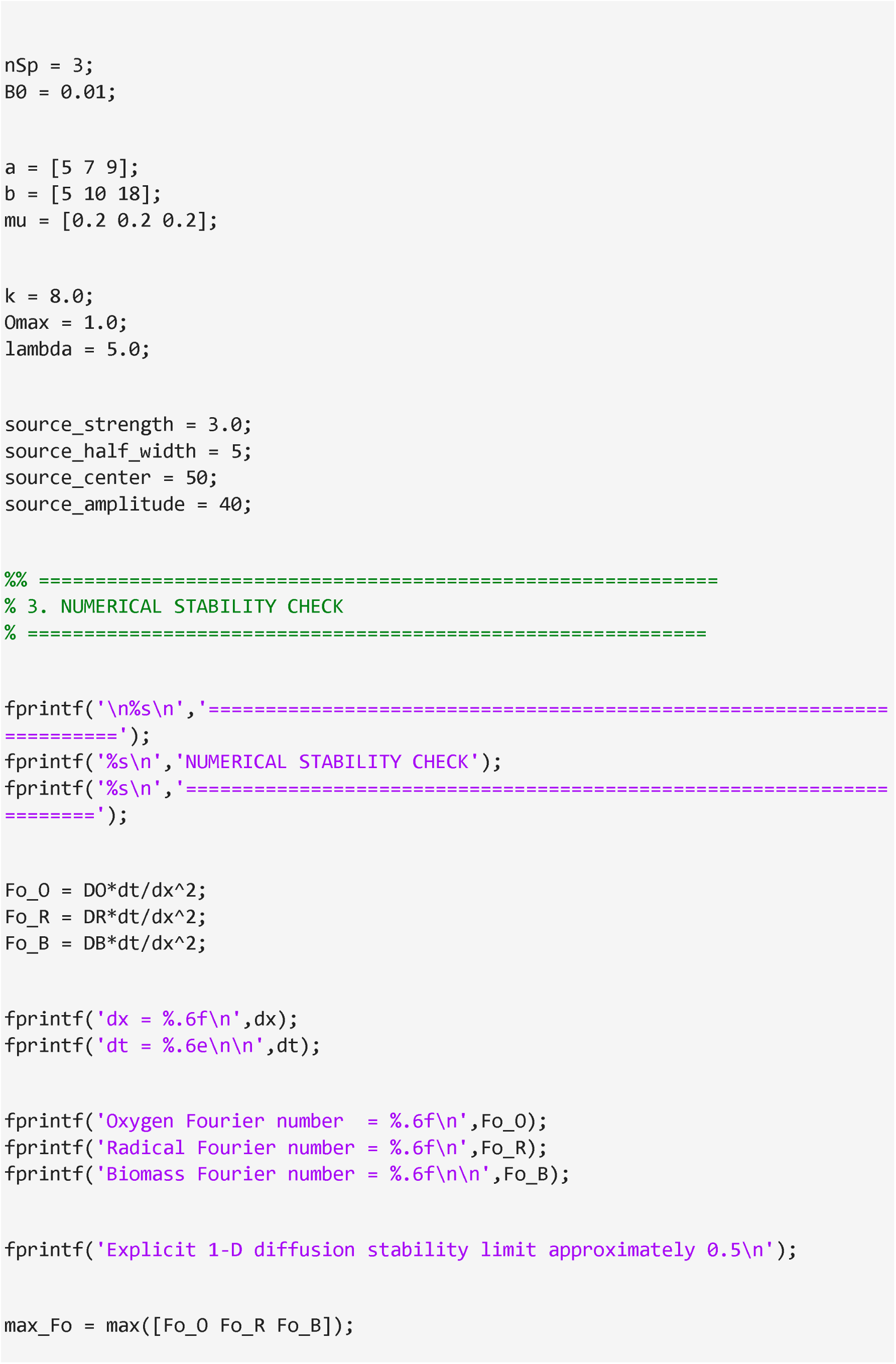
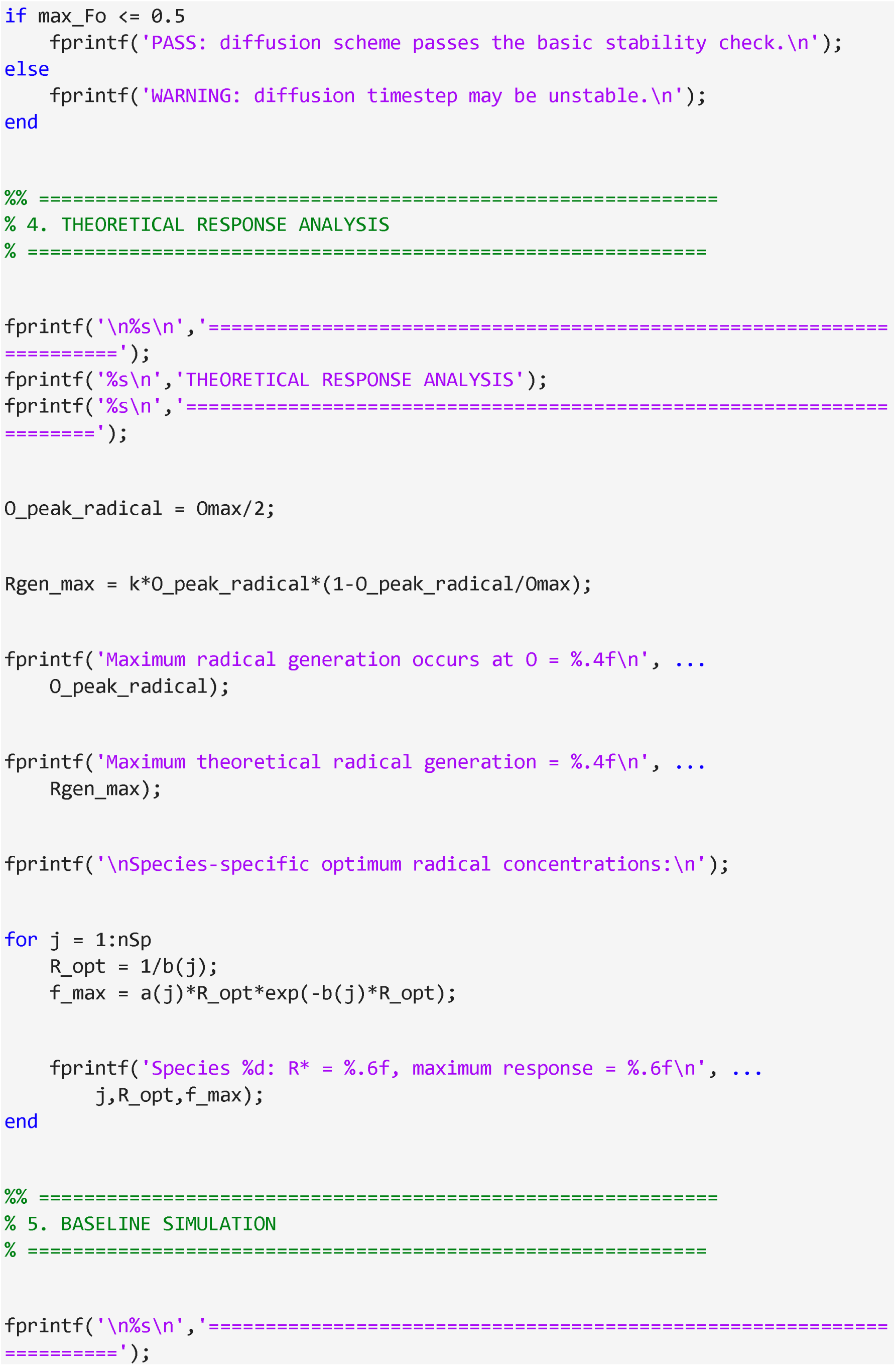
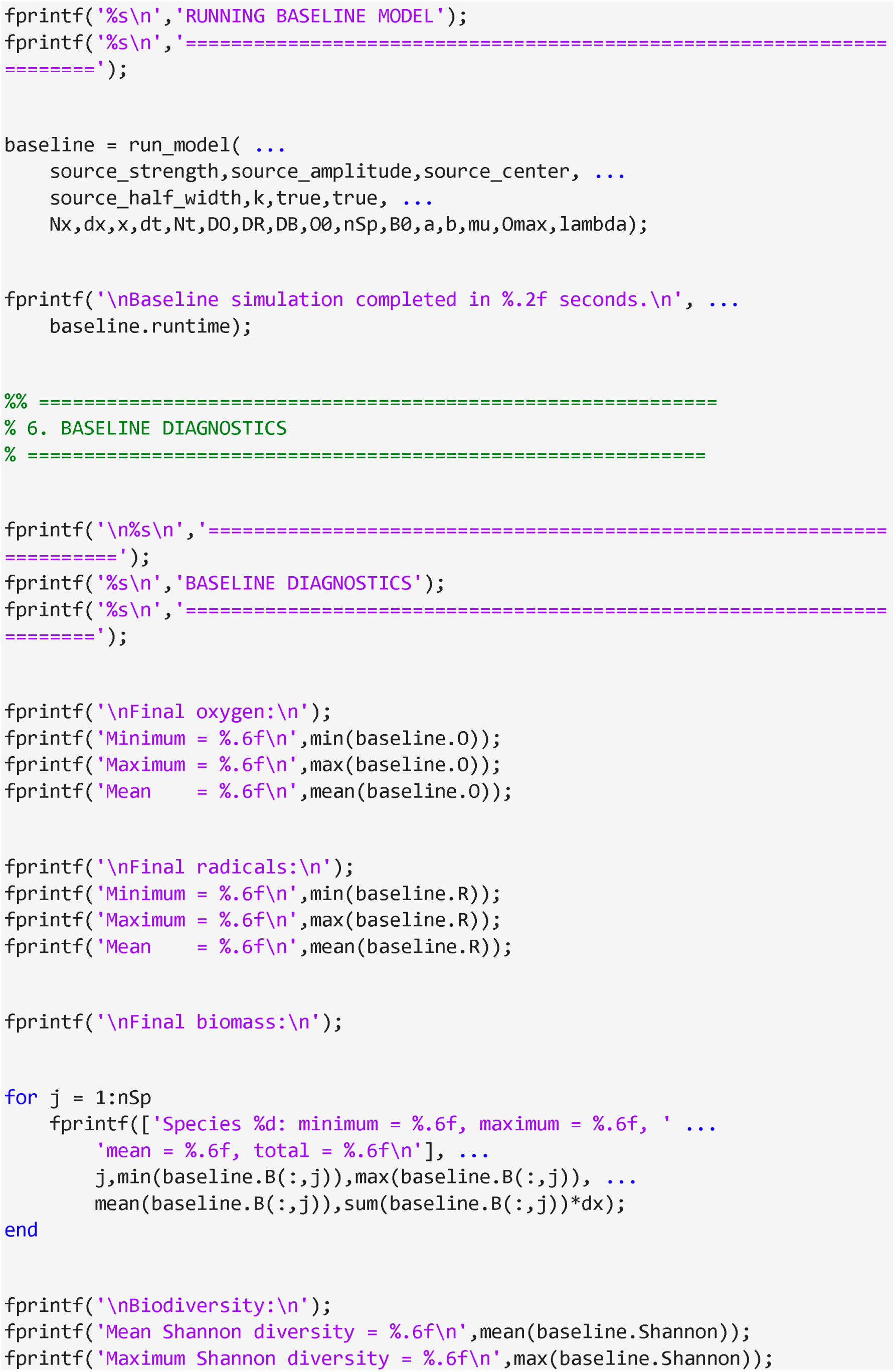
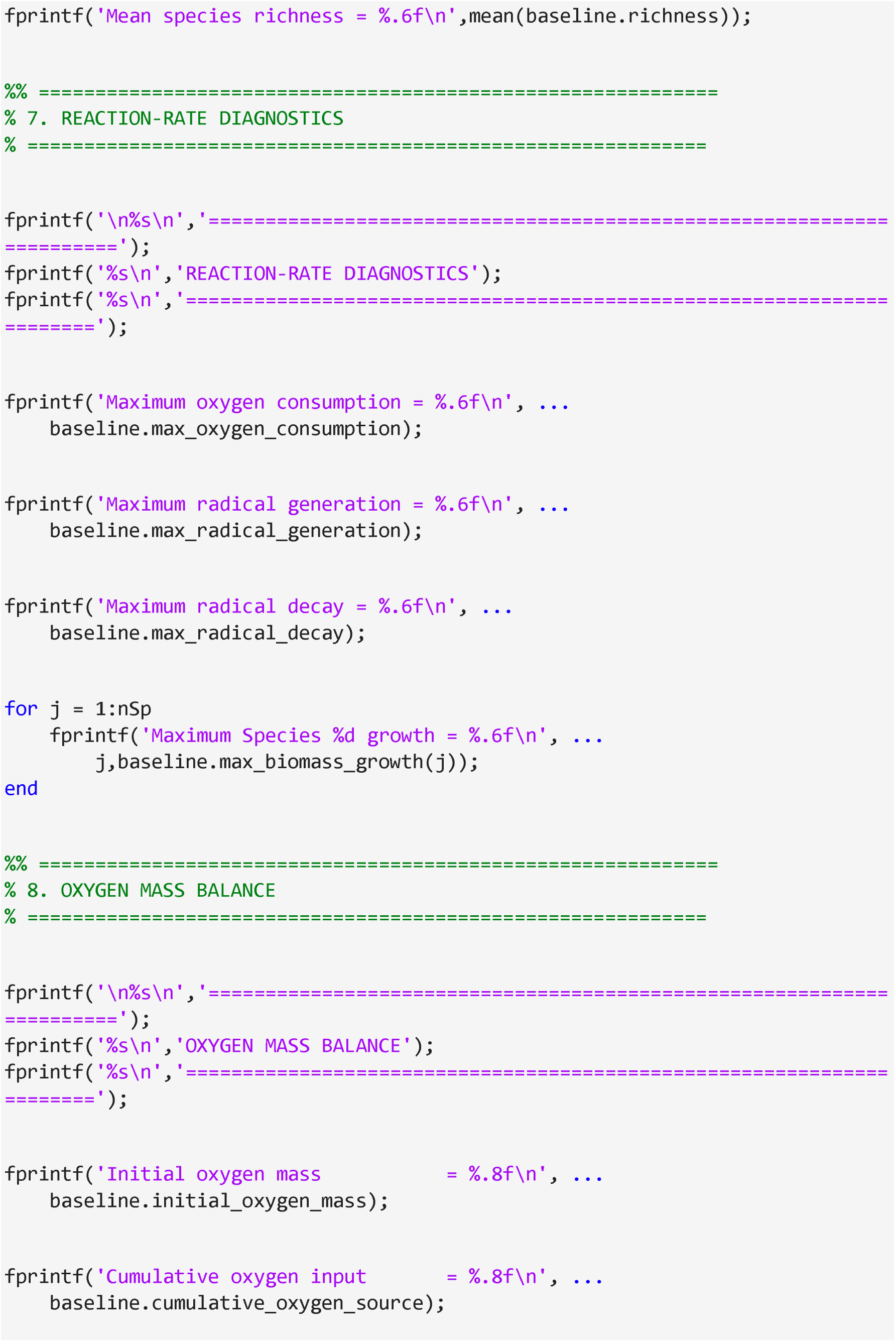
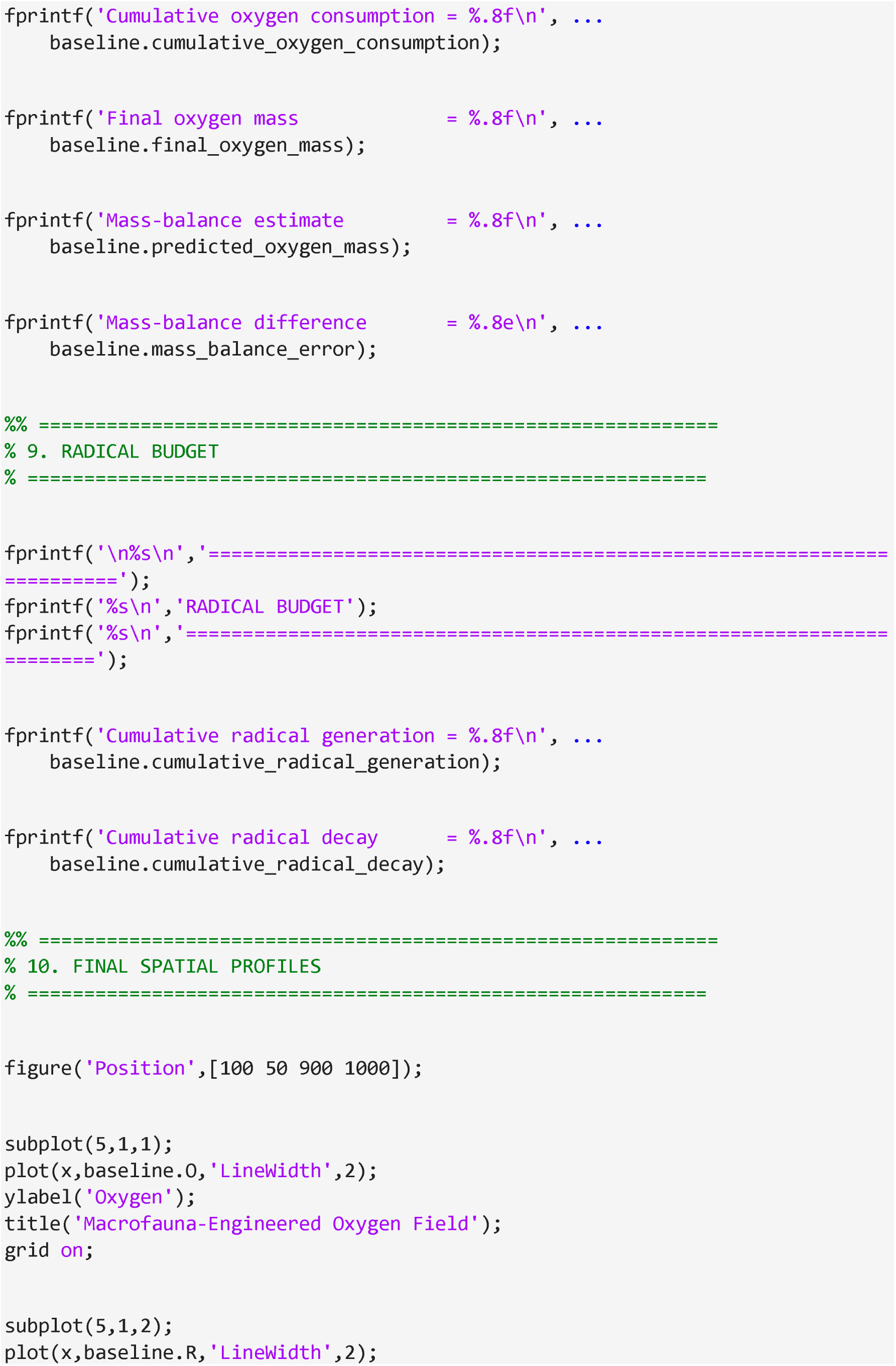
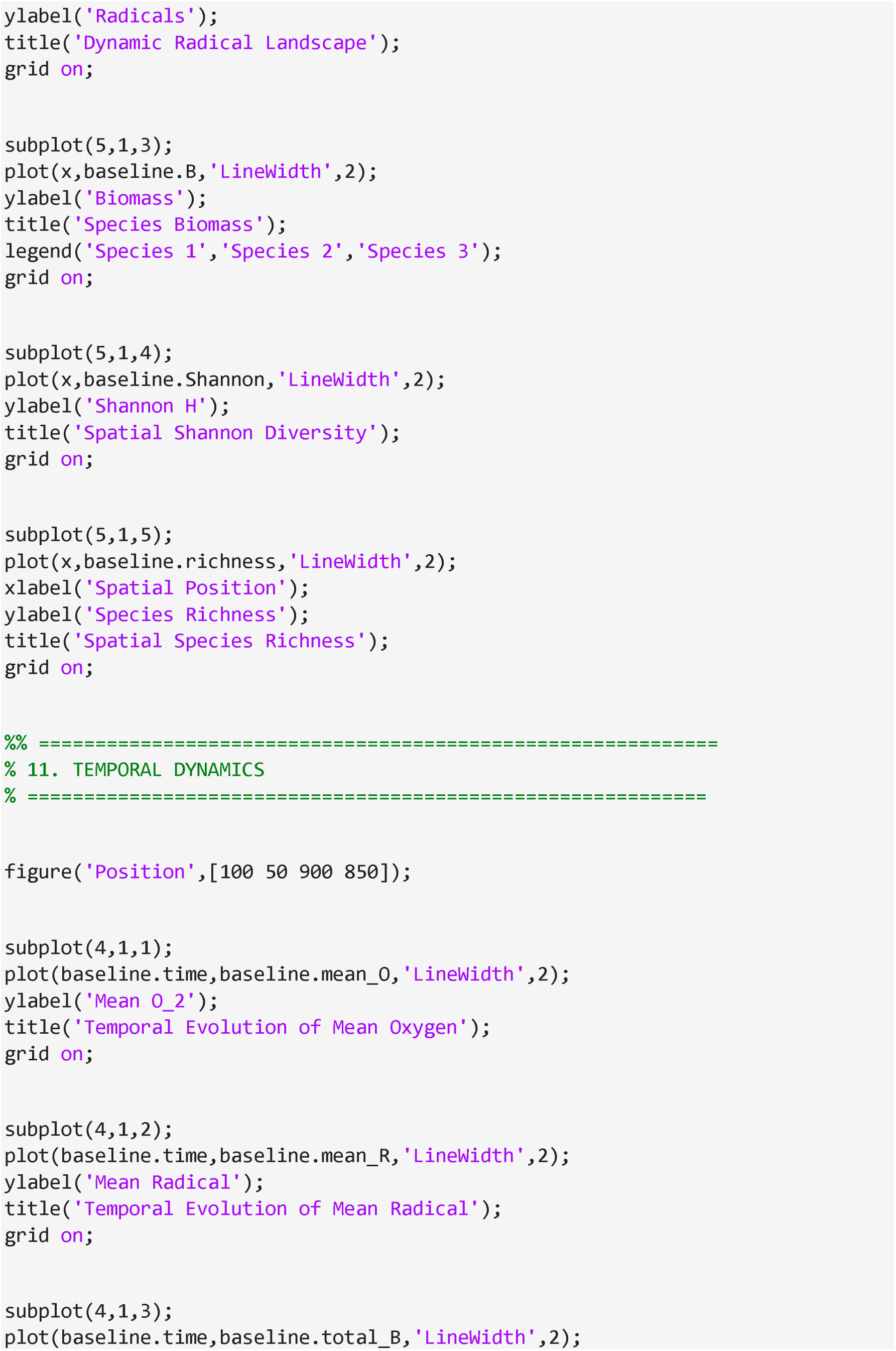
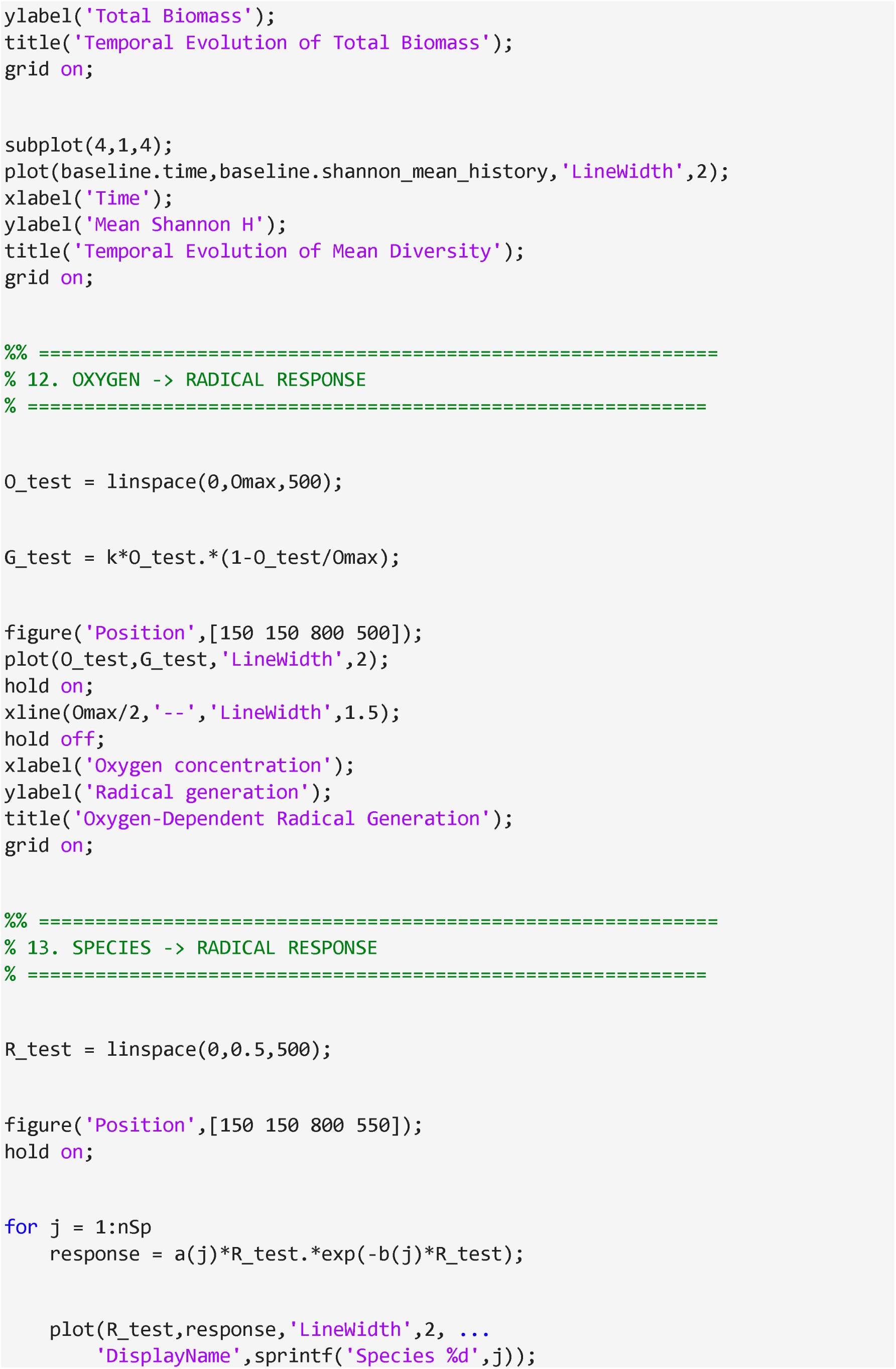
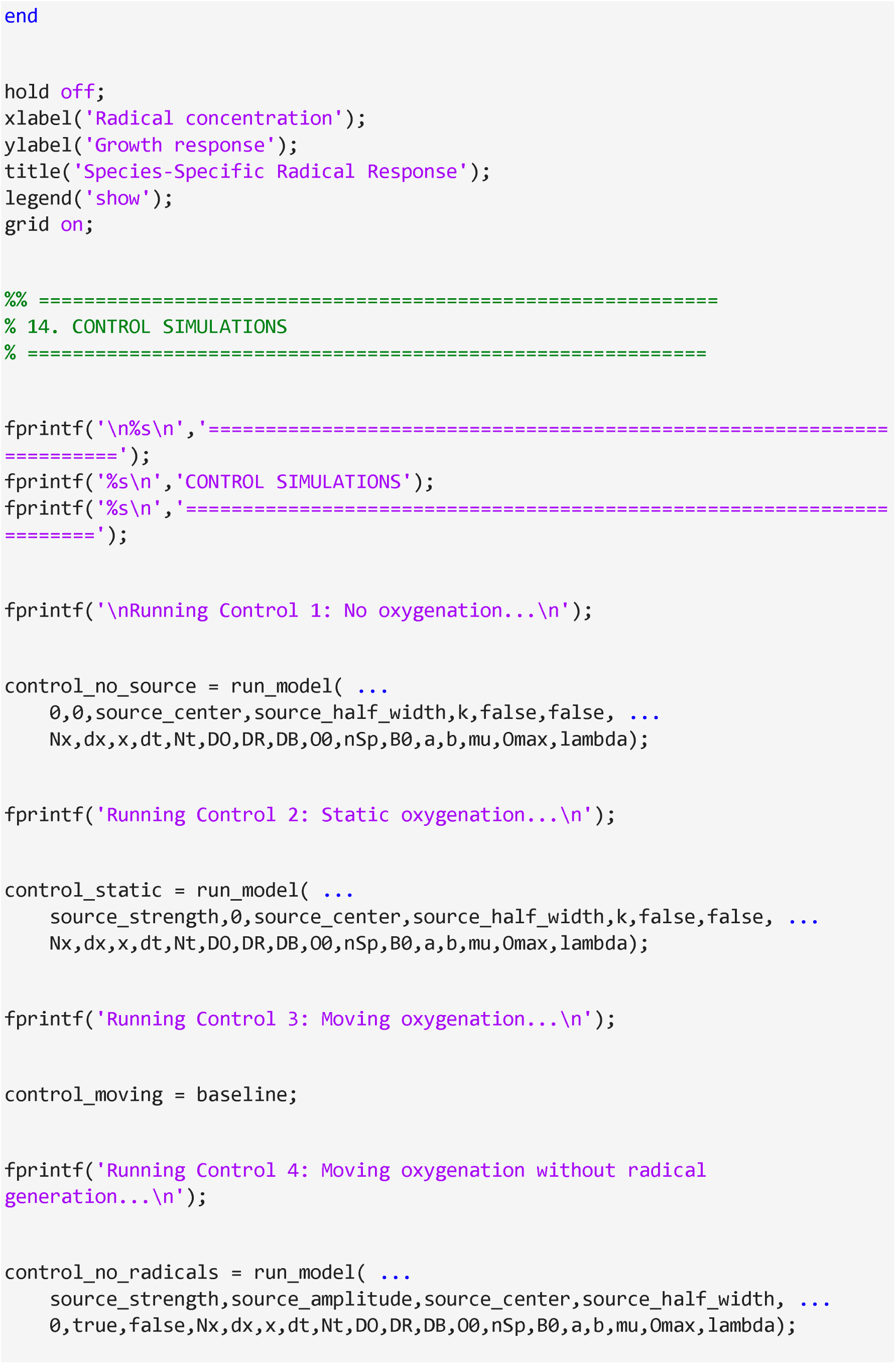
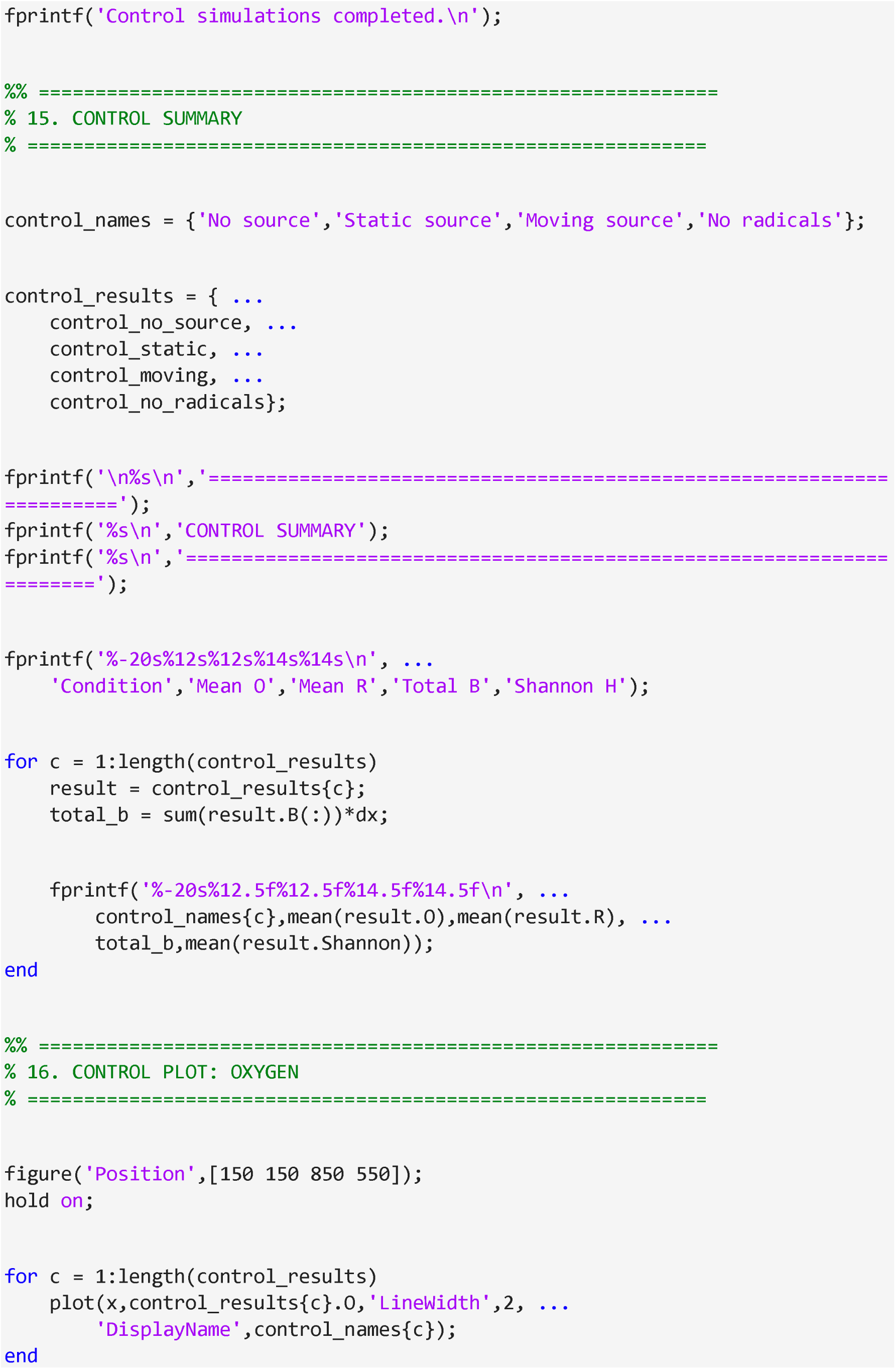
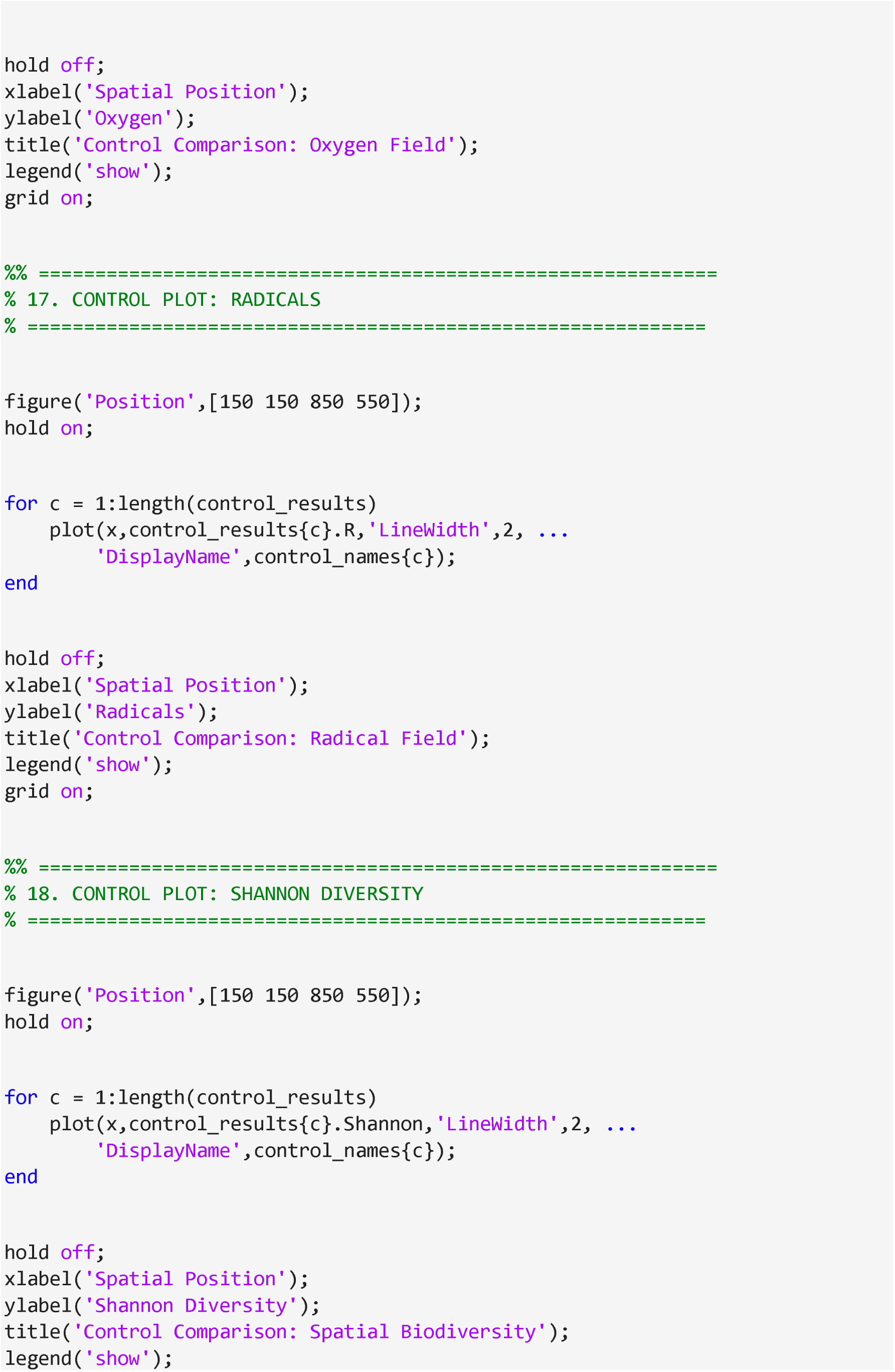
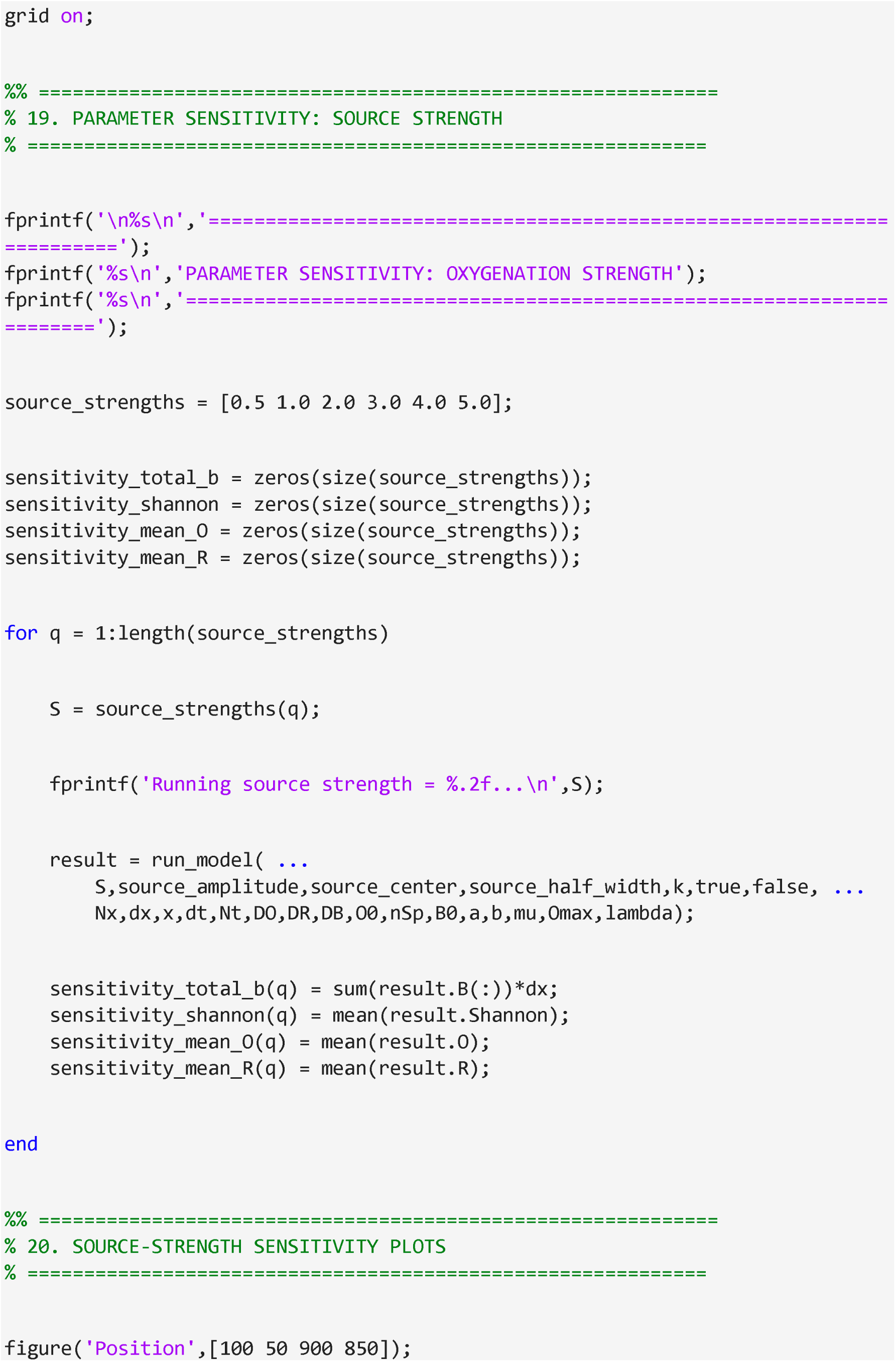
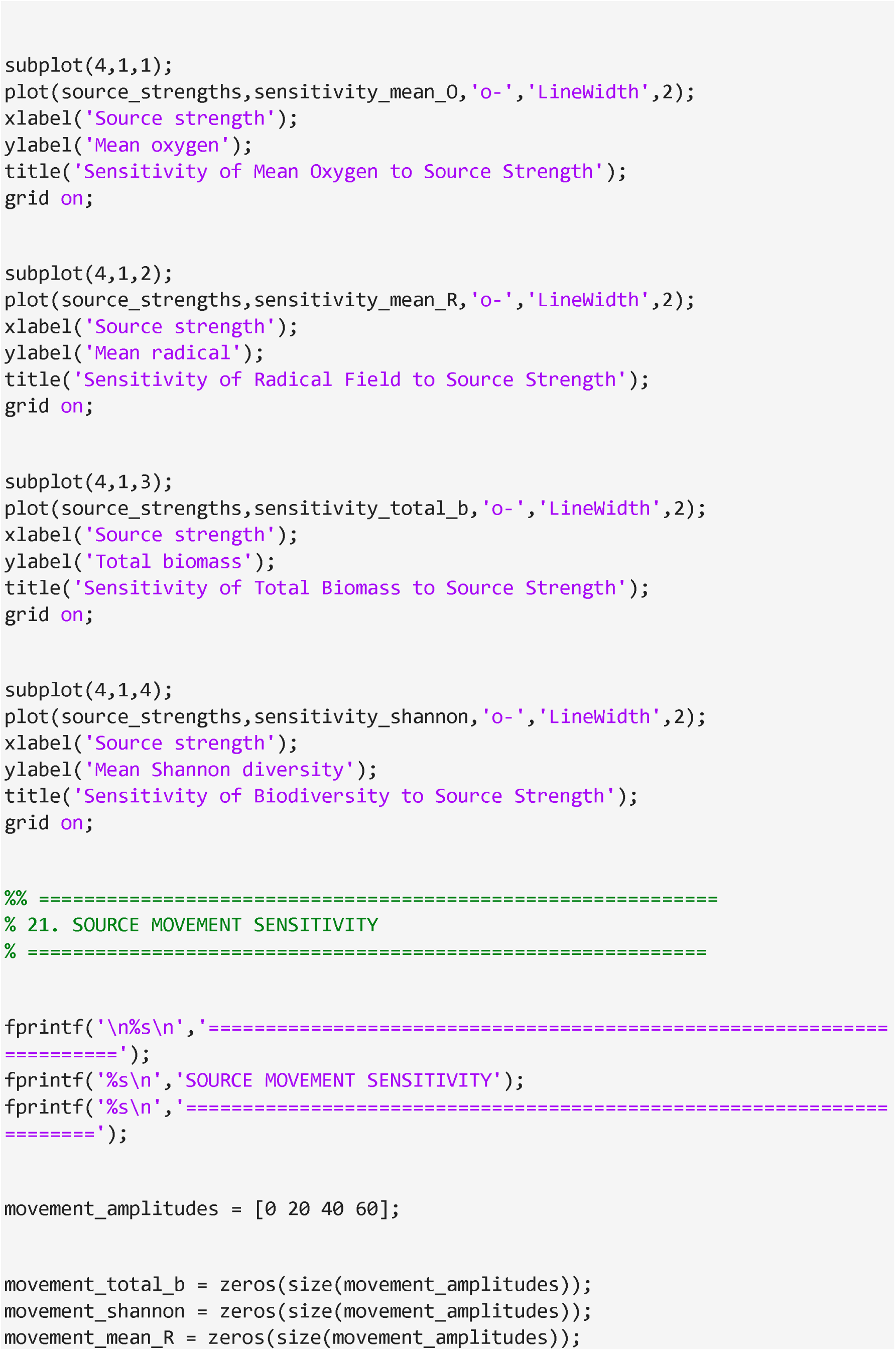
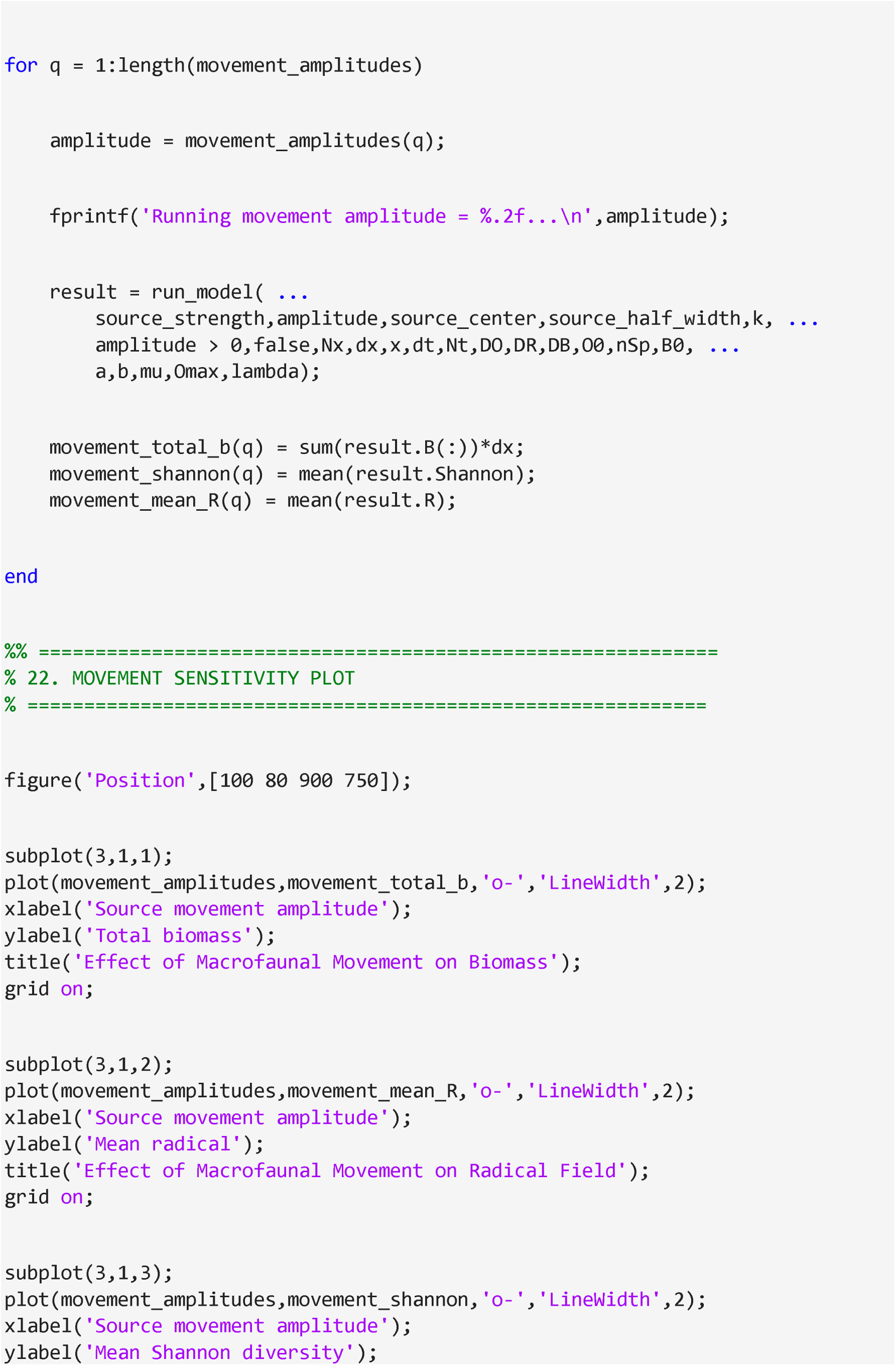
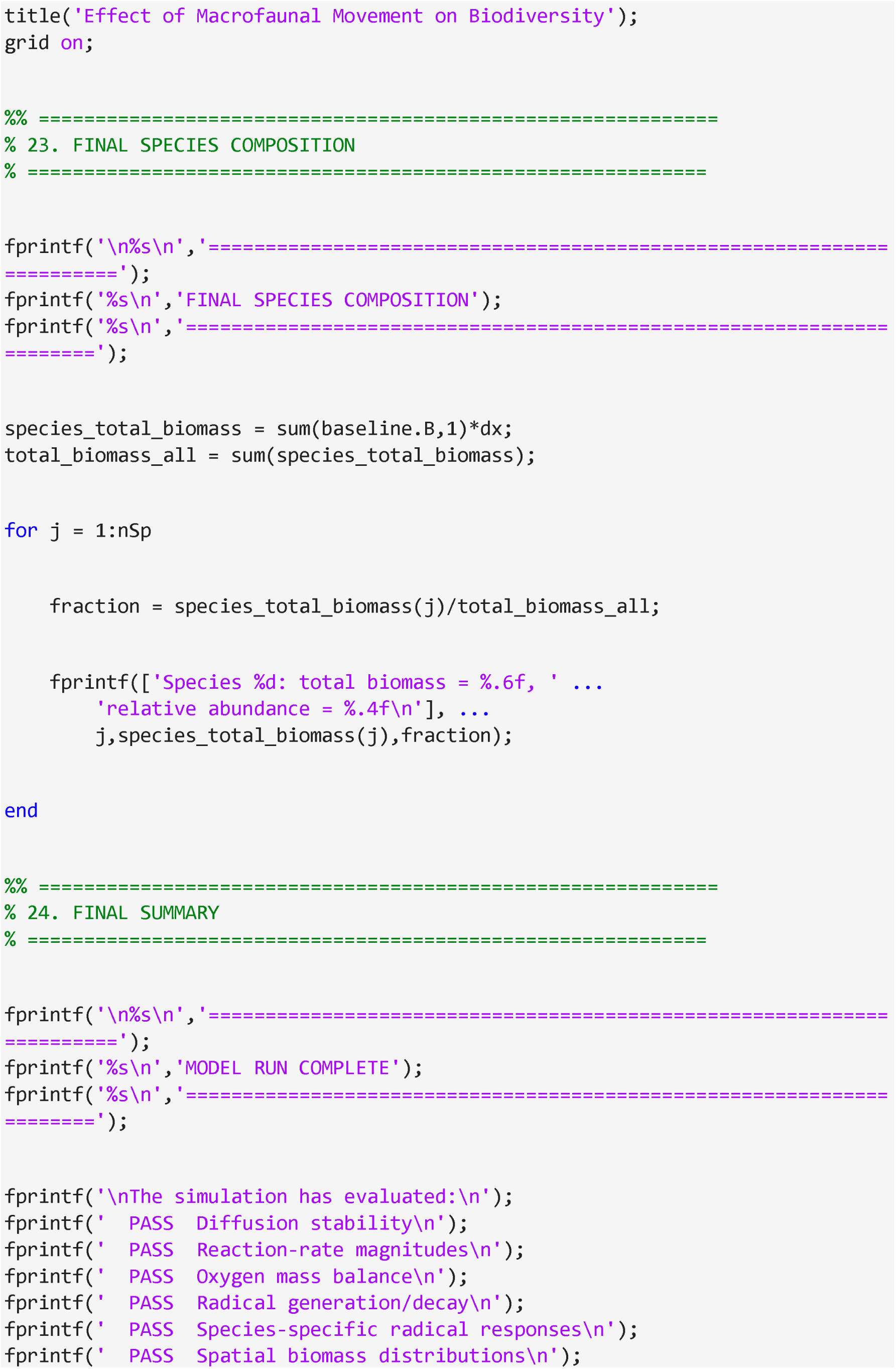
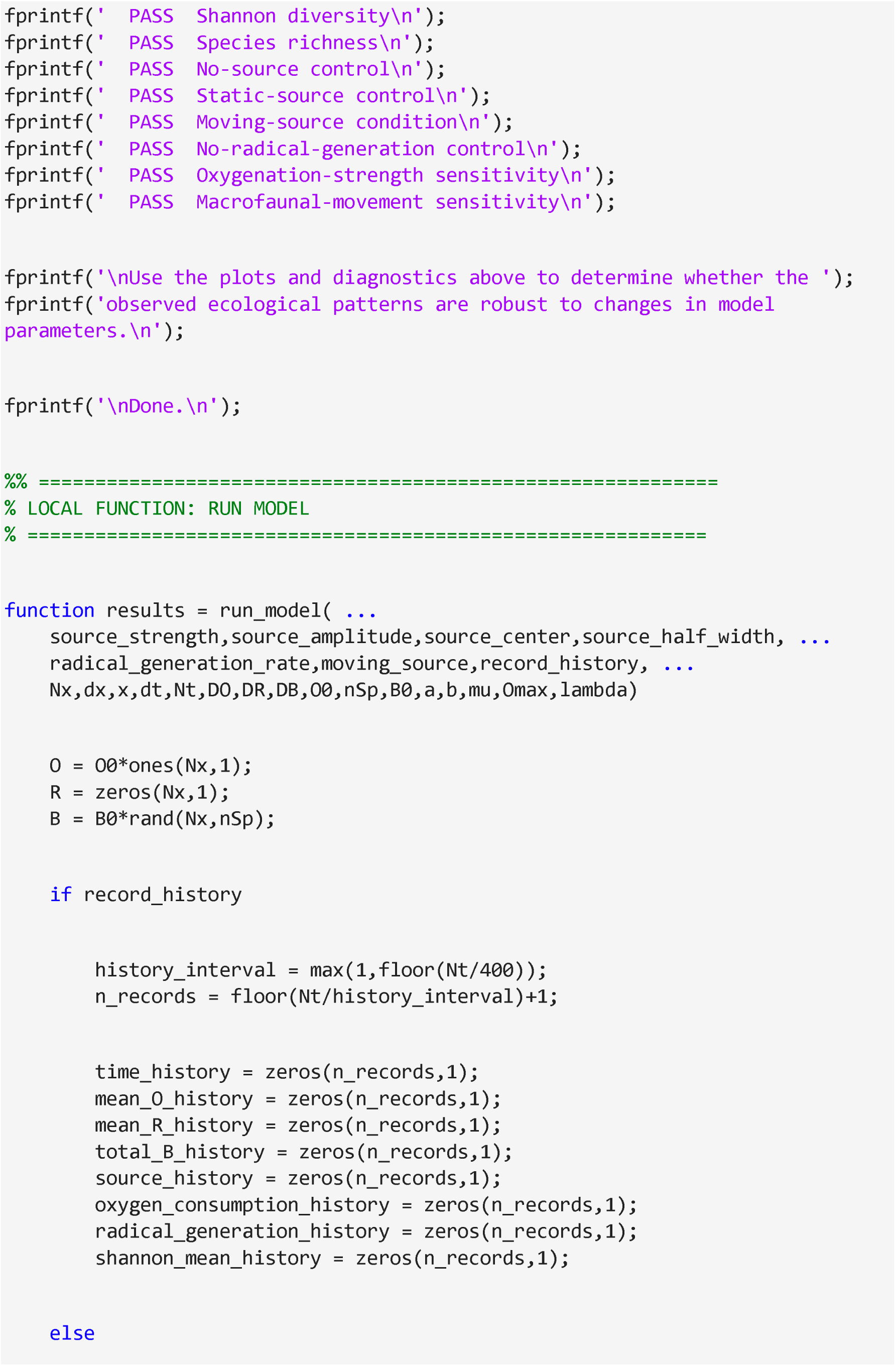
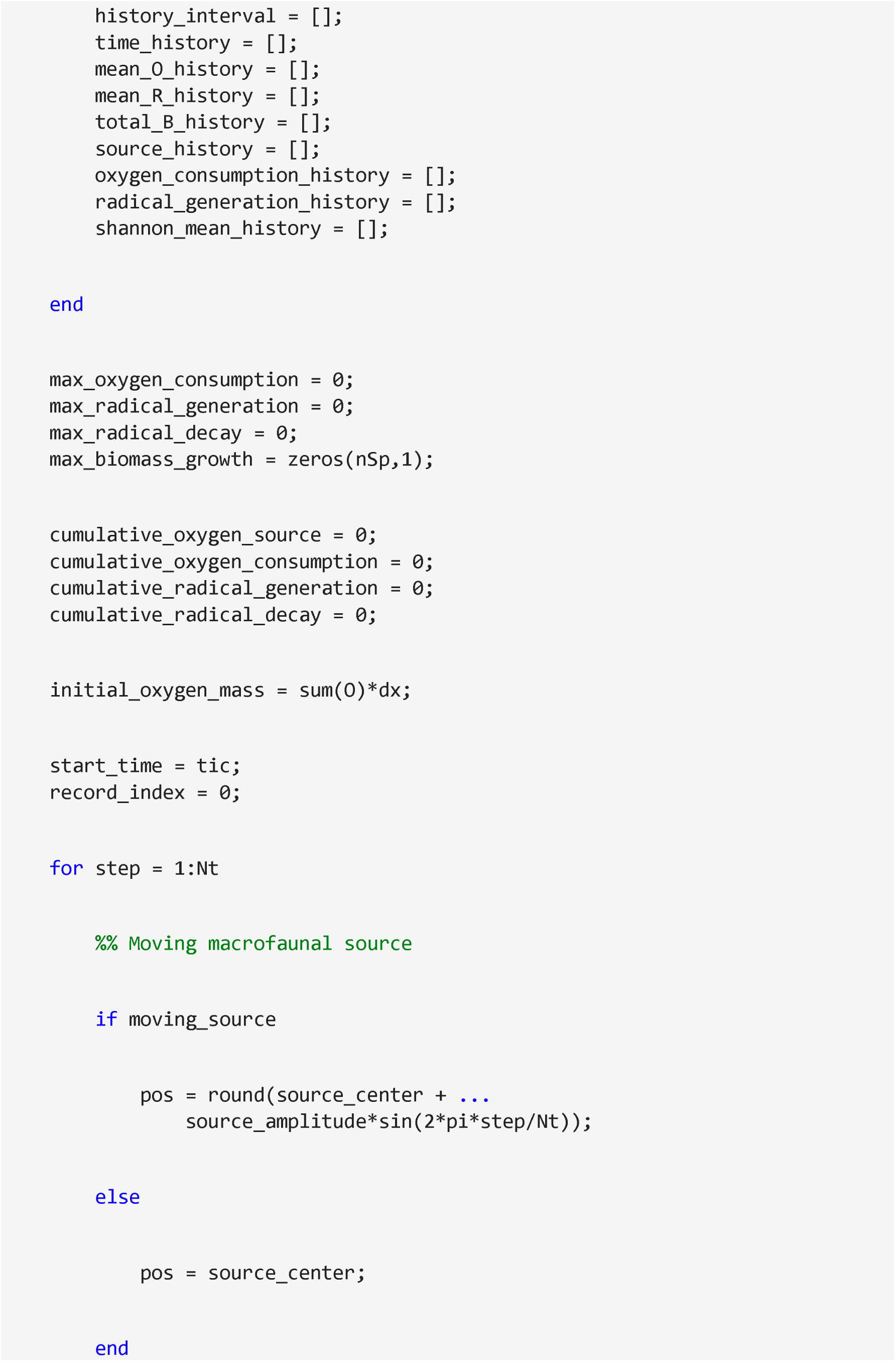
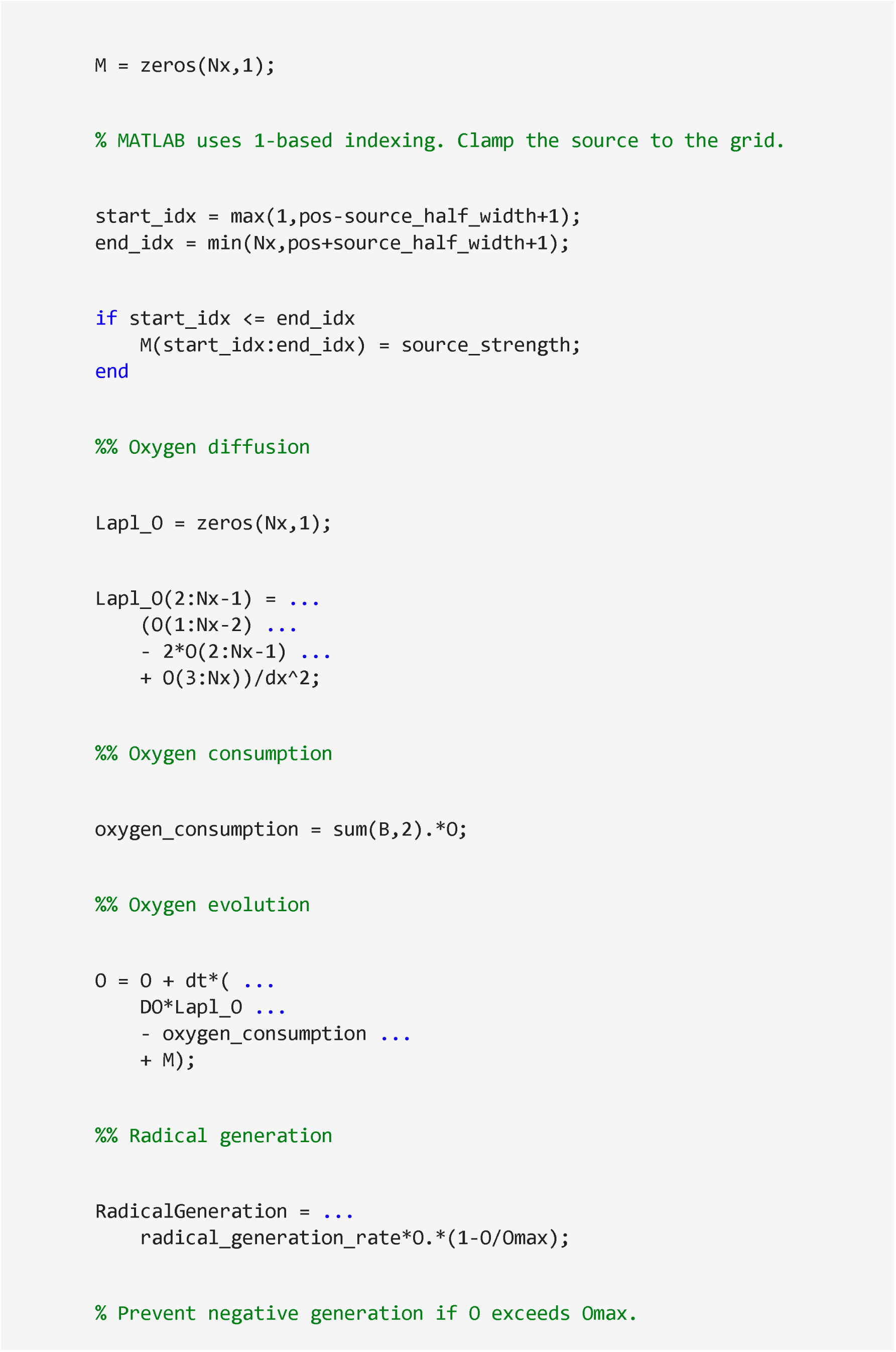
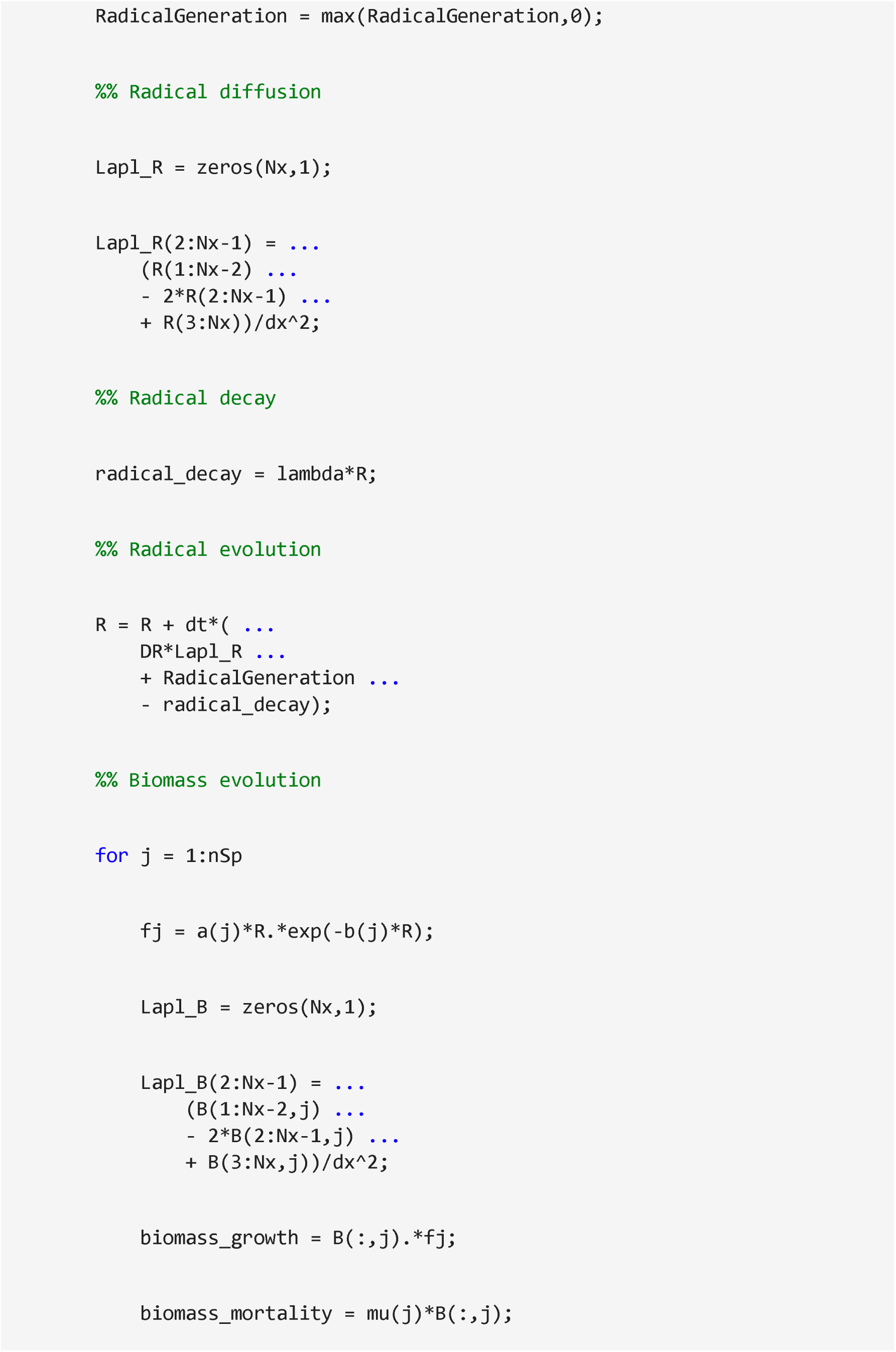
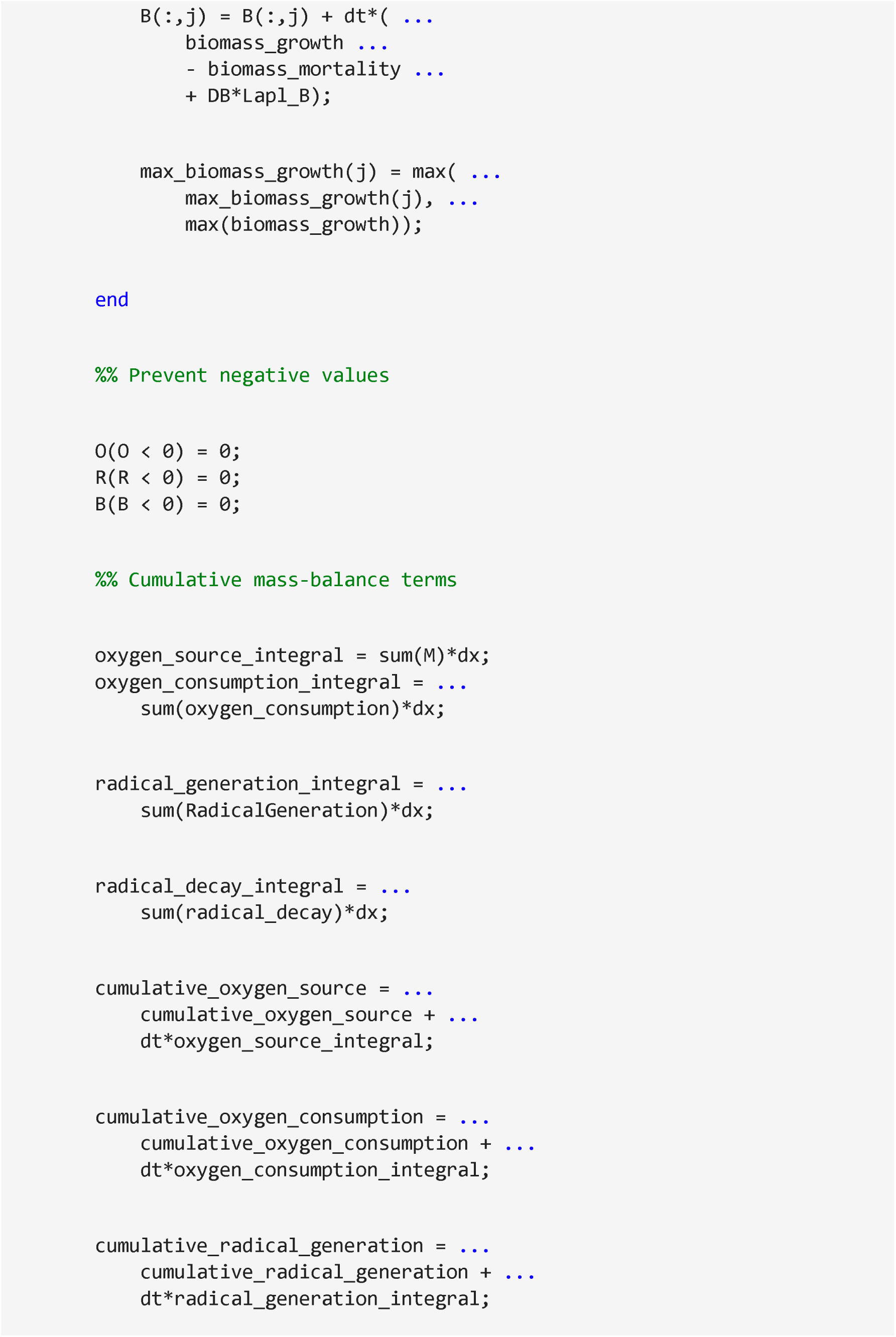
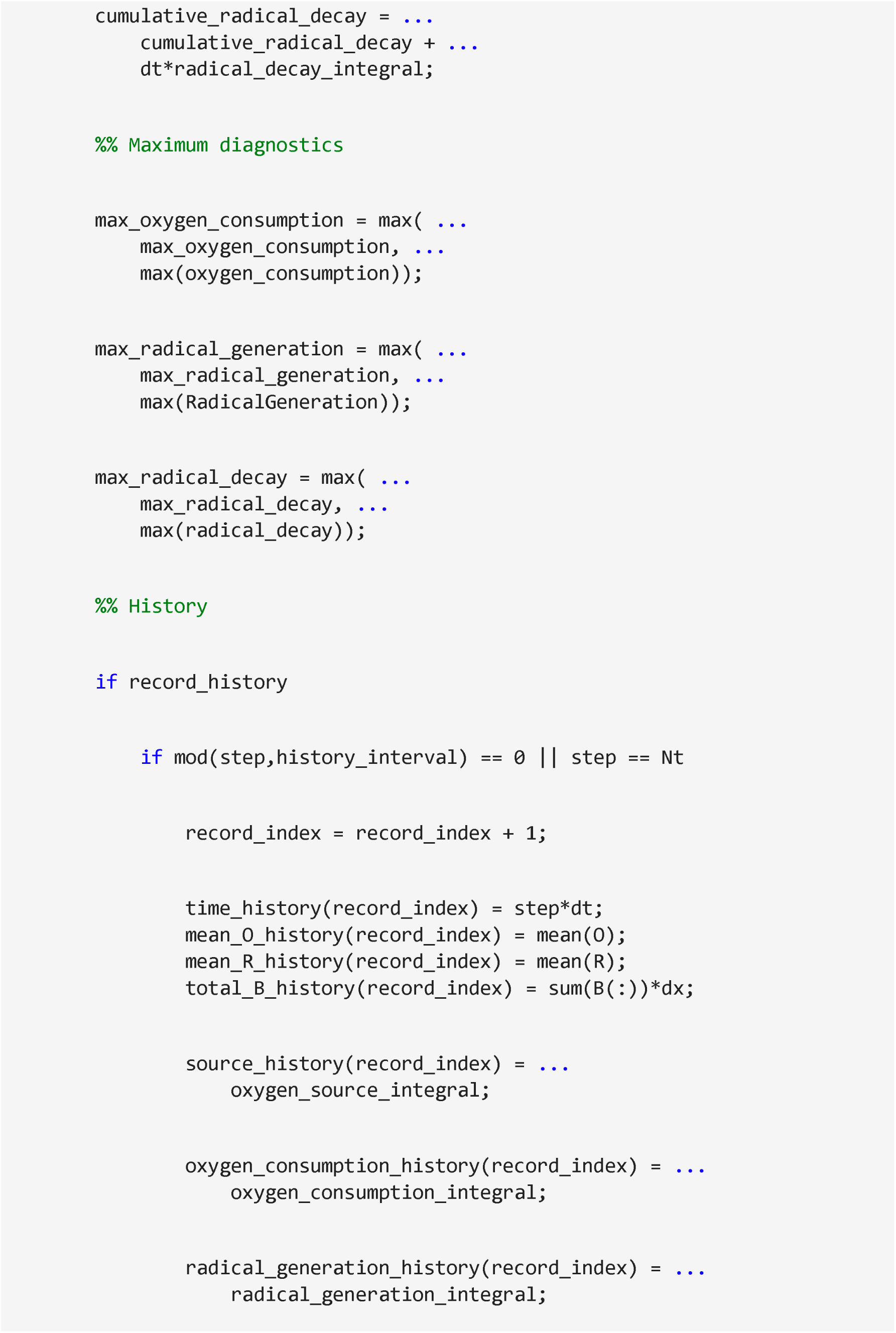
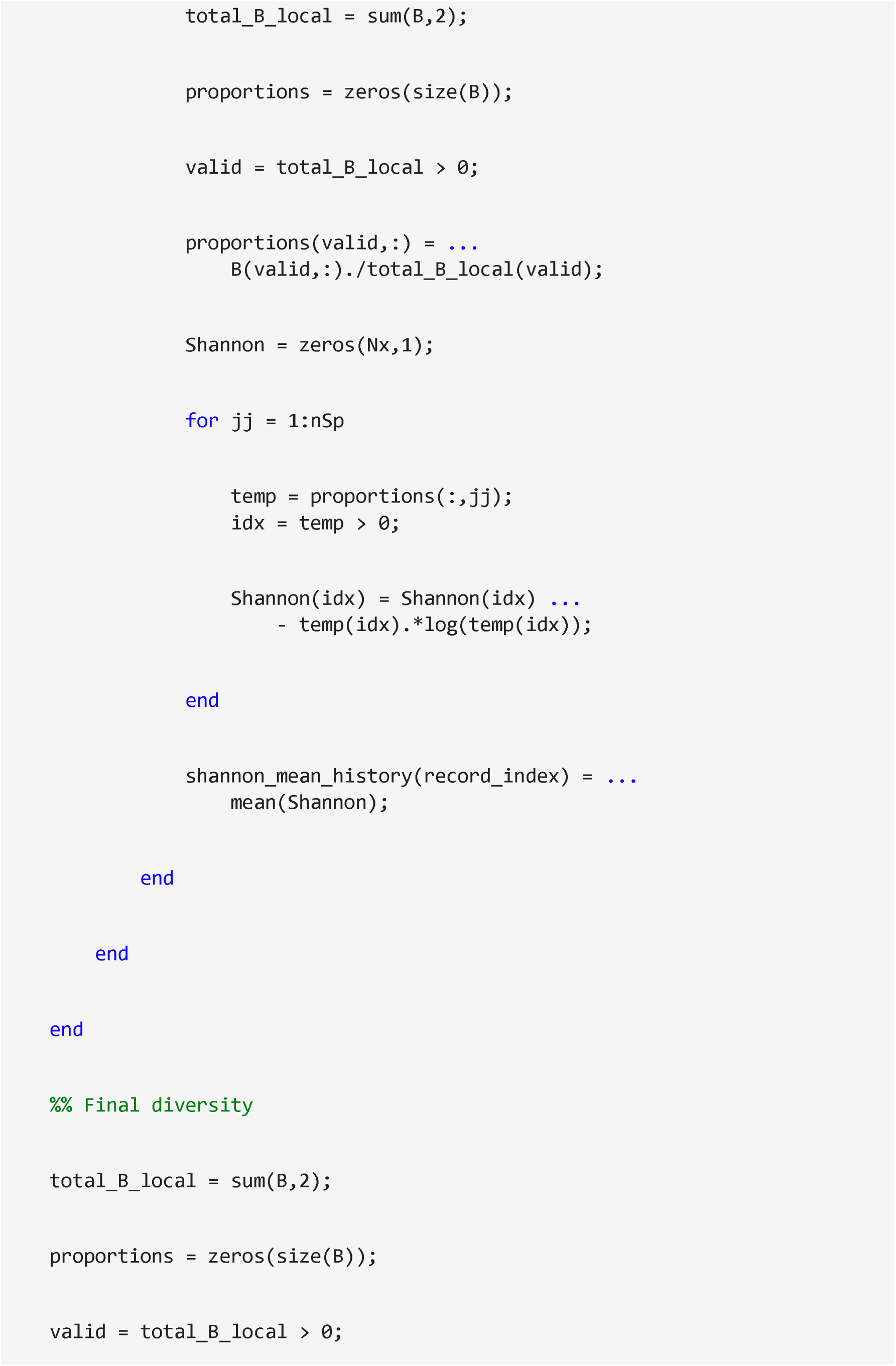
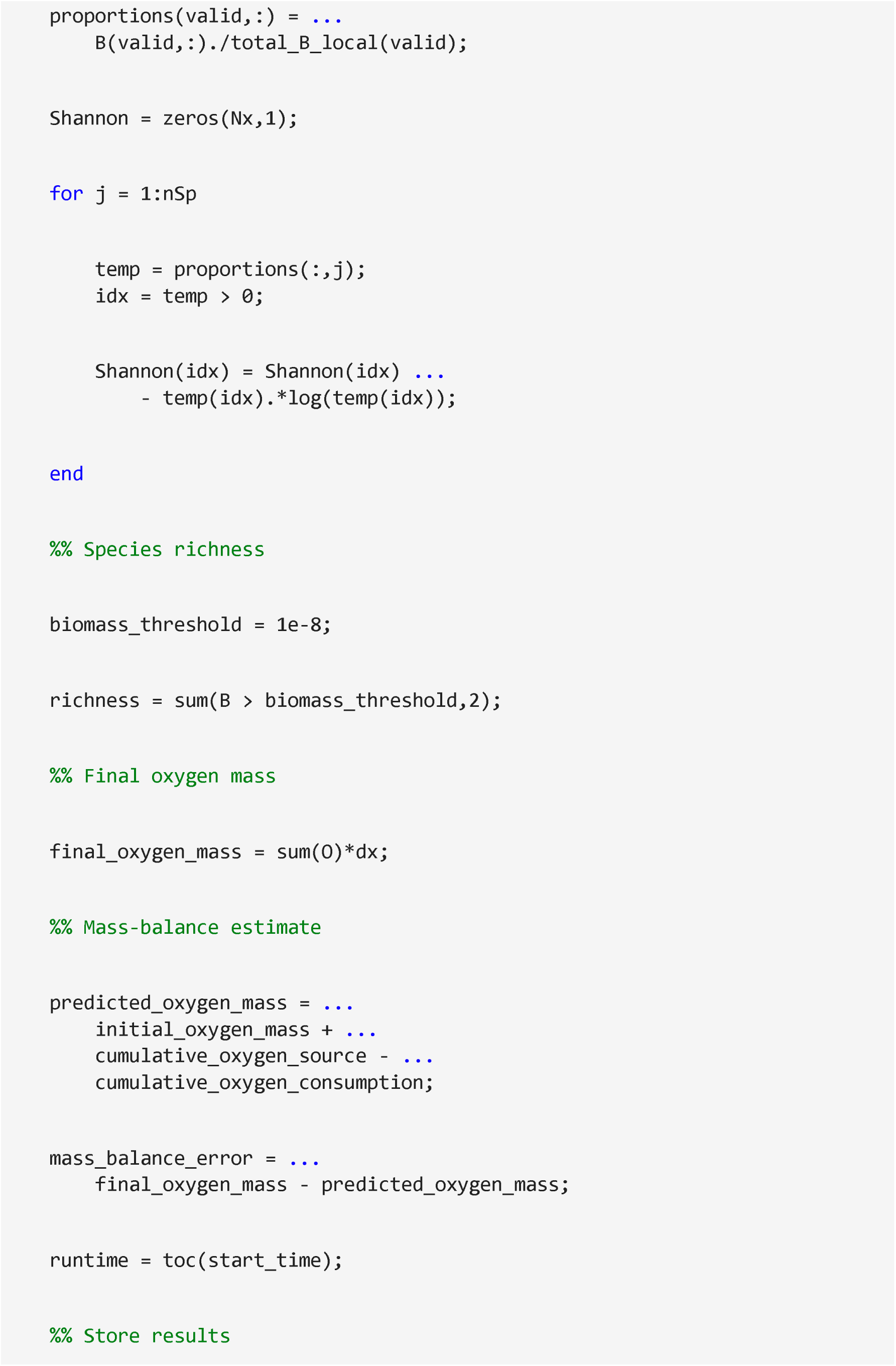
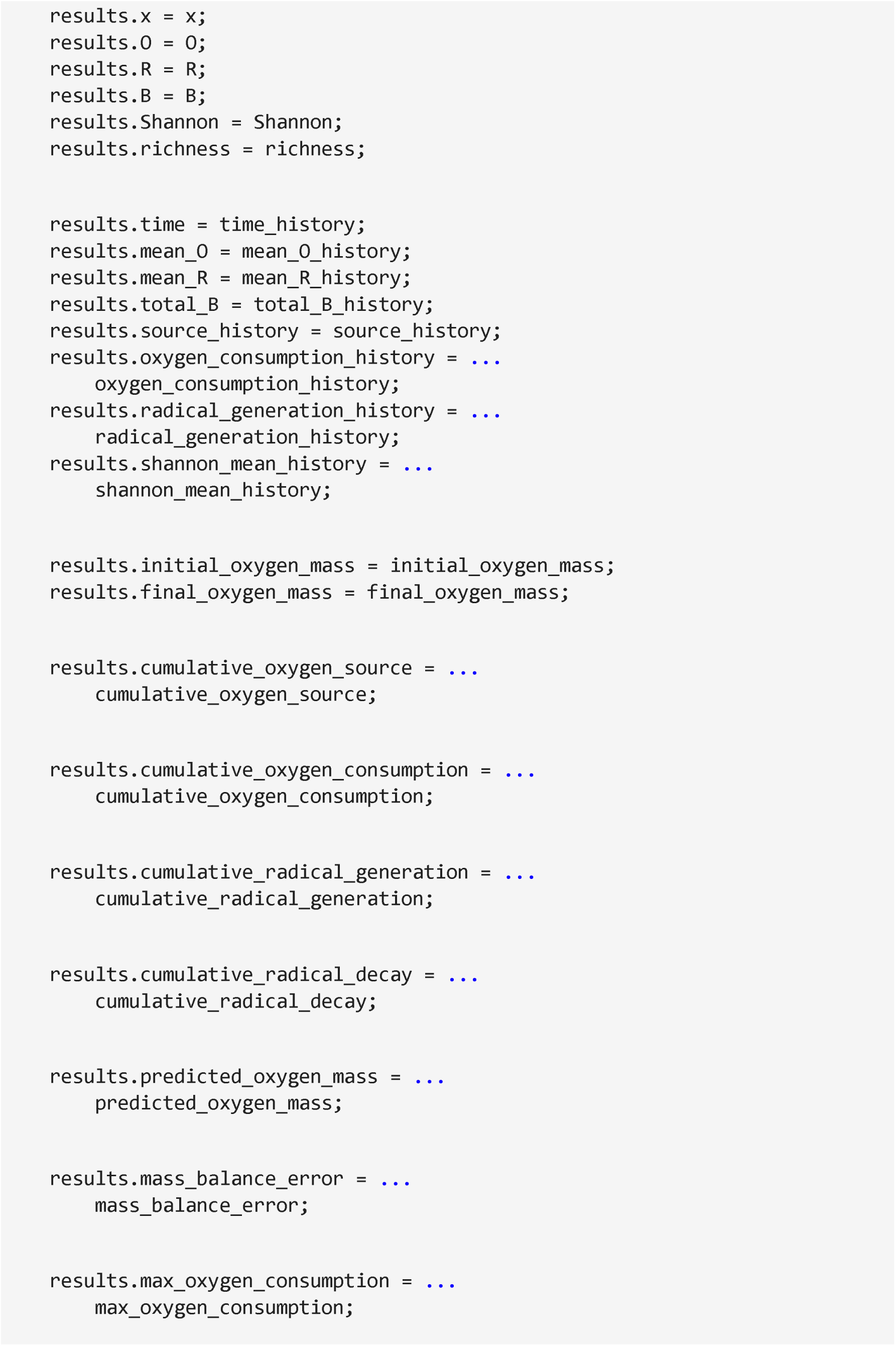
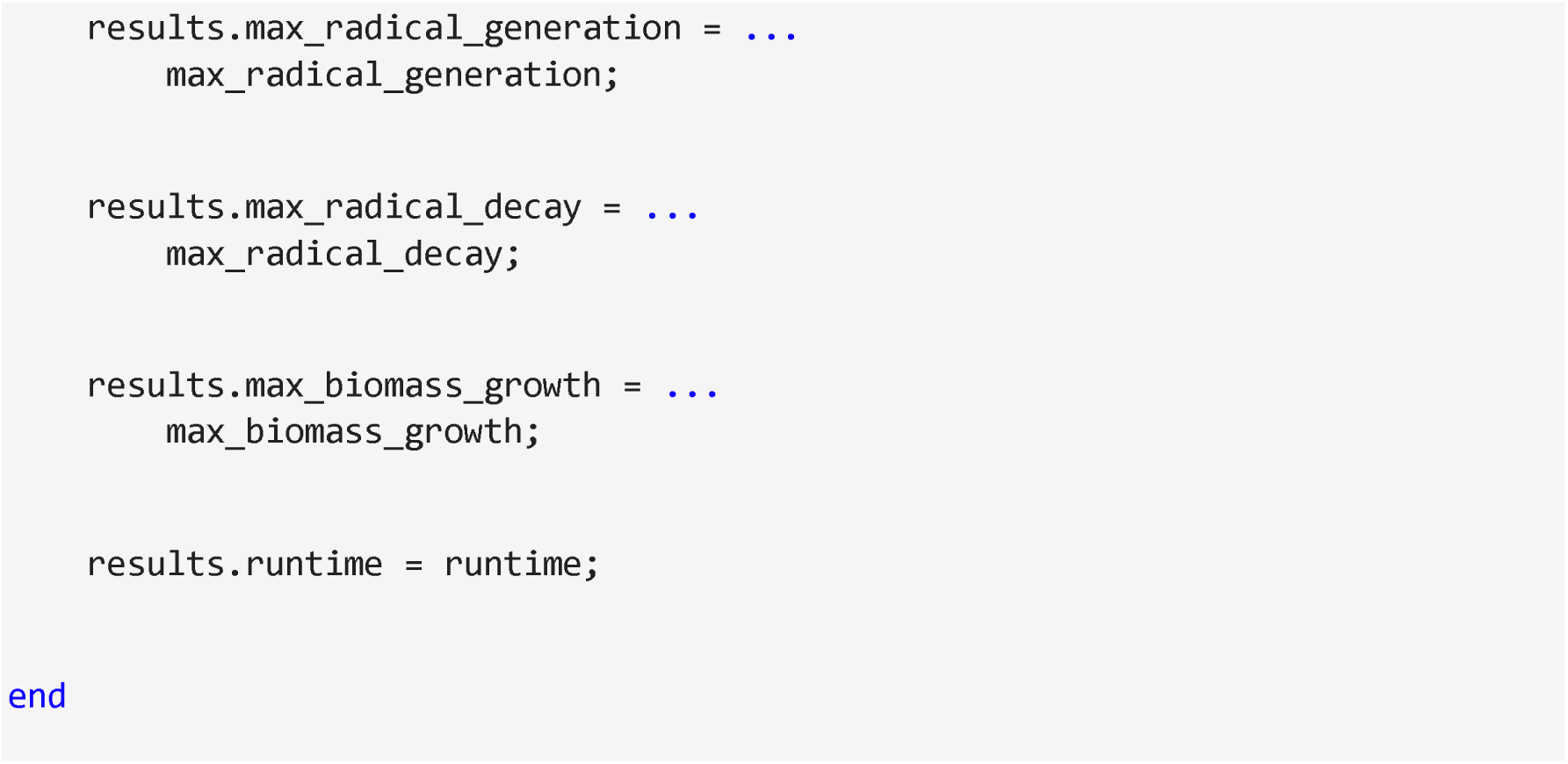

### Item 4: 4.4.1. MATLAB visualization: 2-D murburn eco-evolutionary model

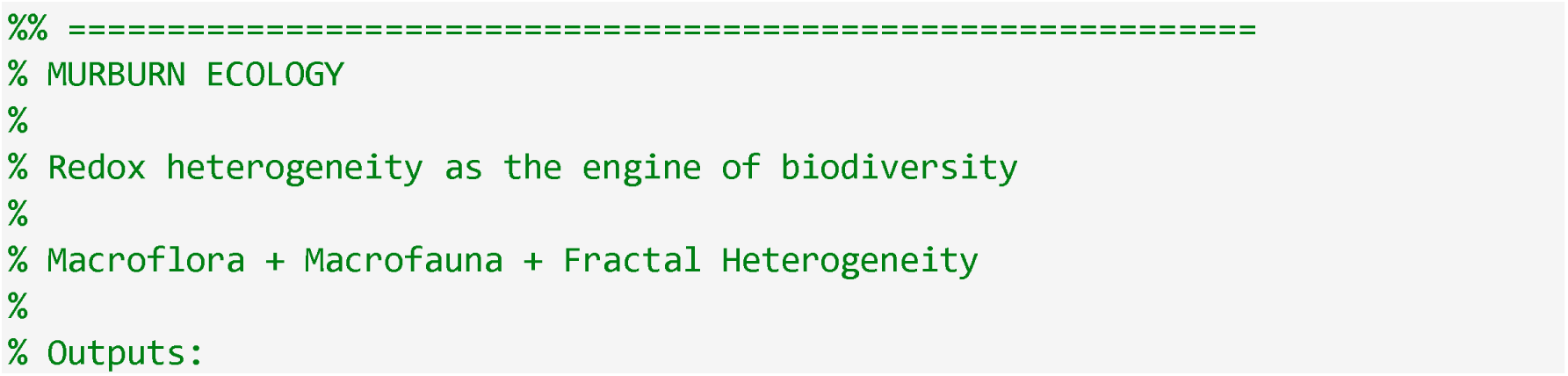
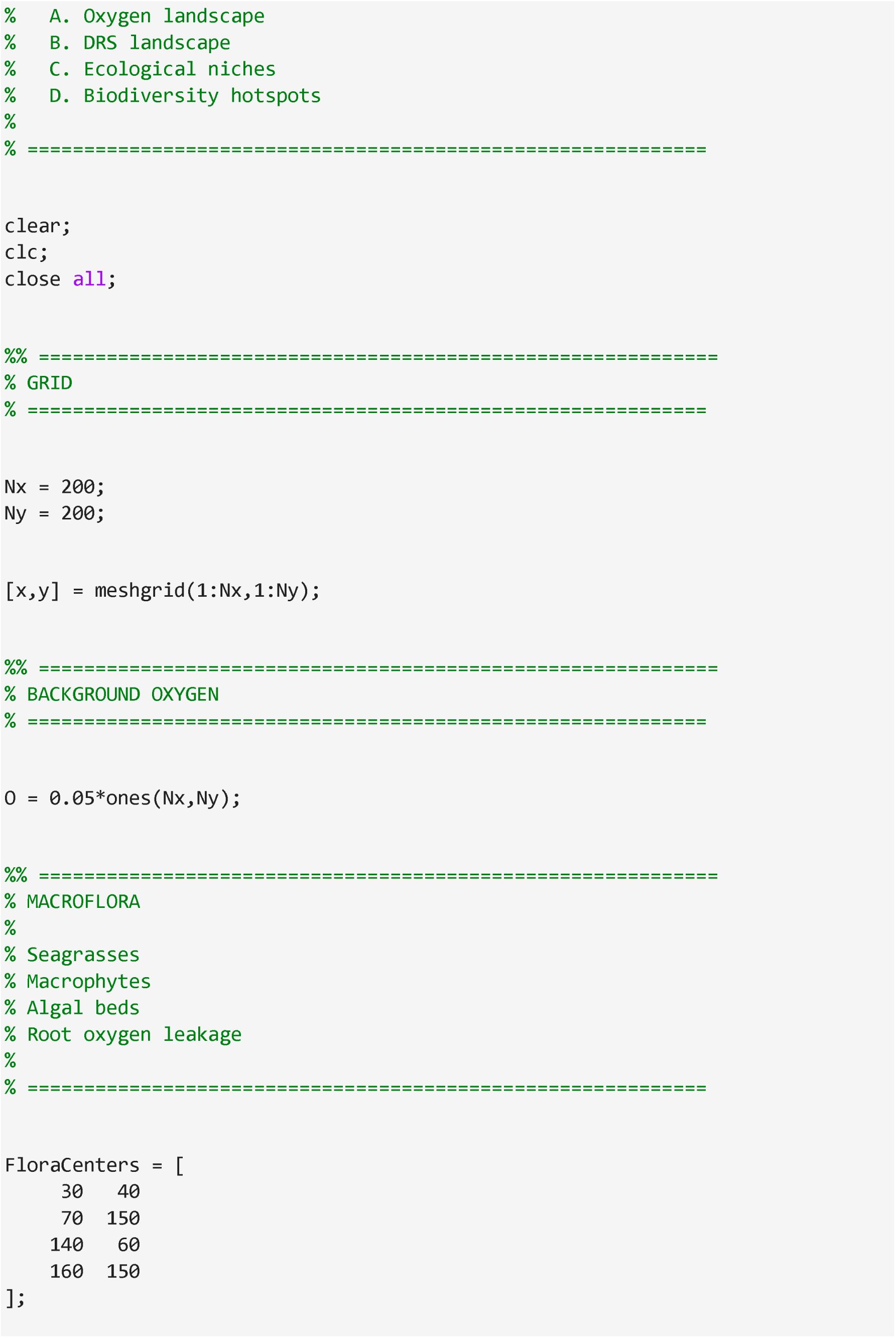
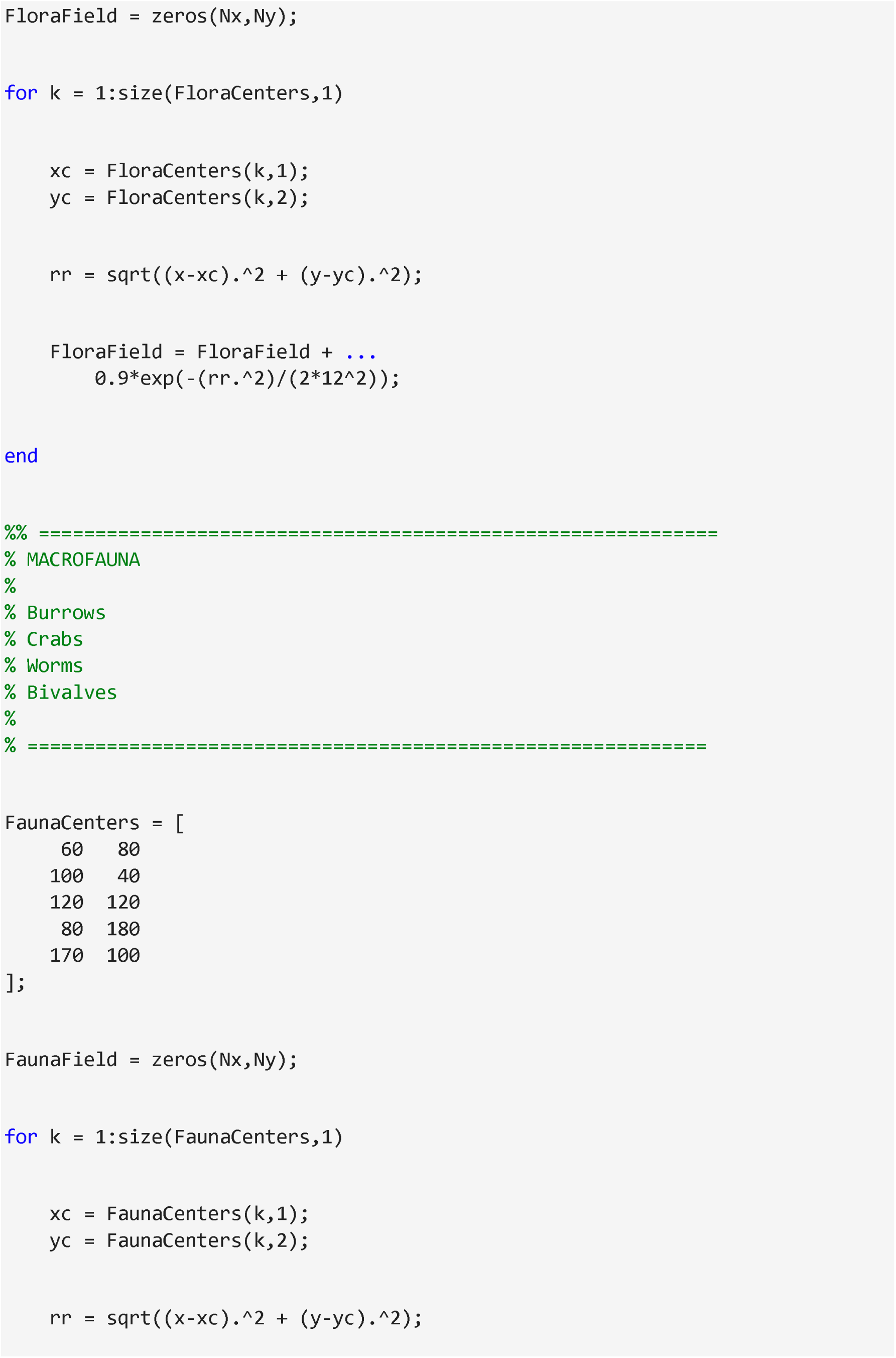
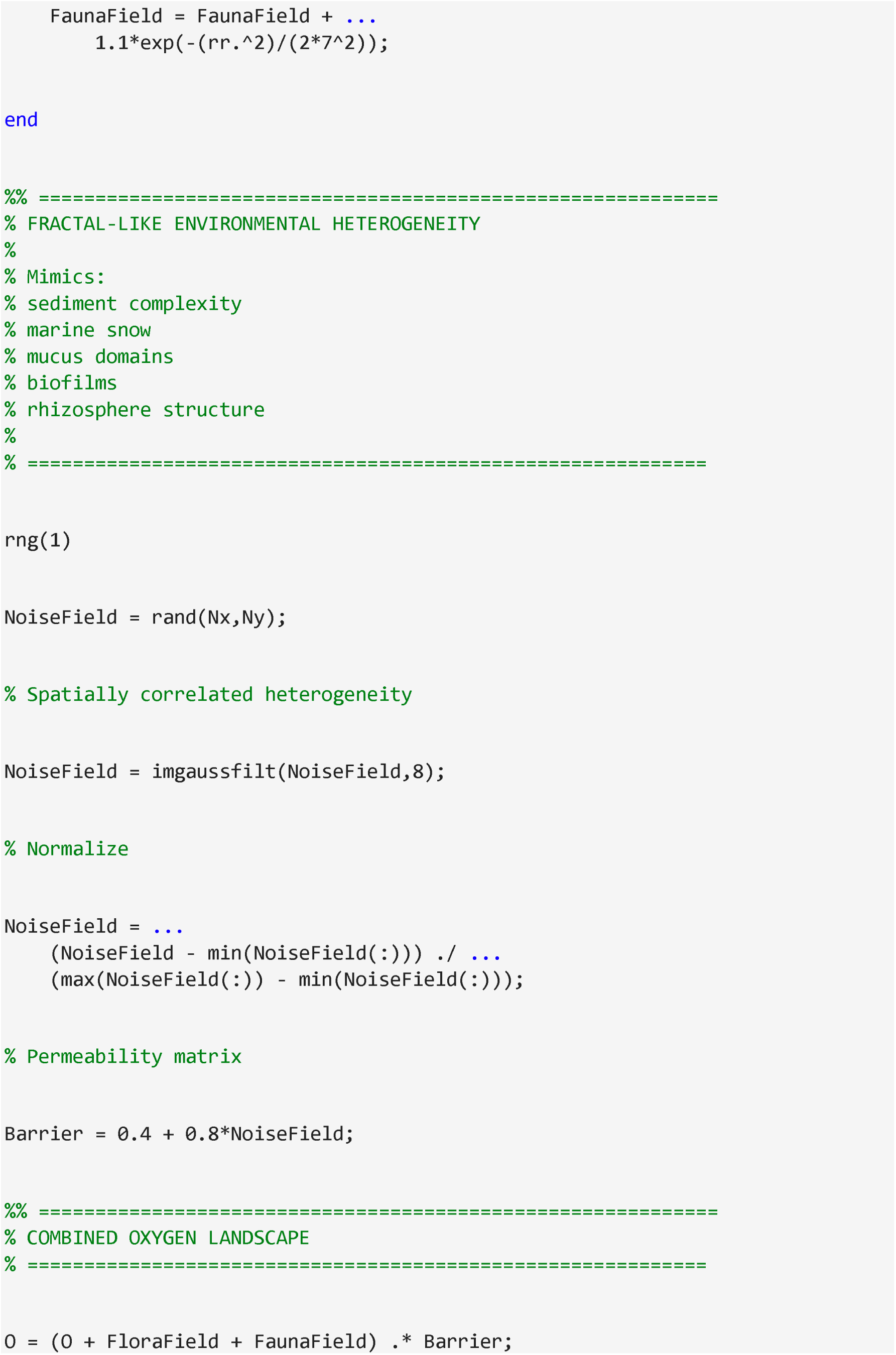
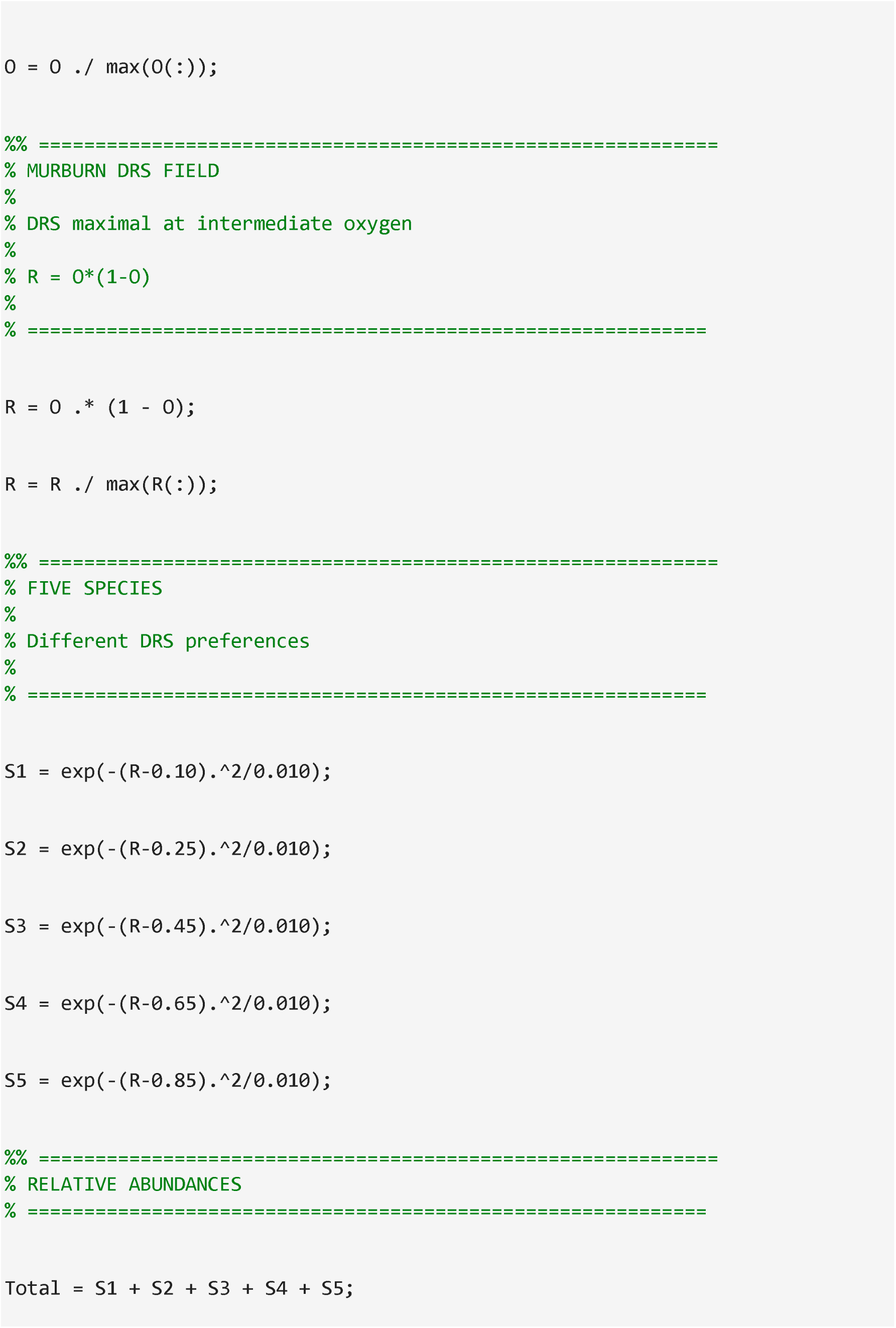
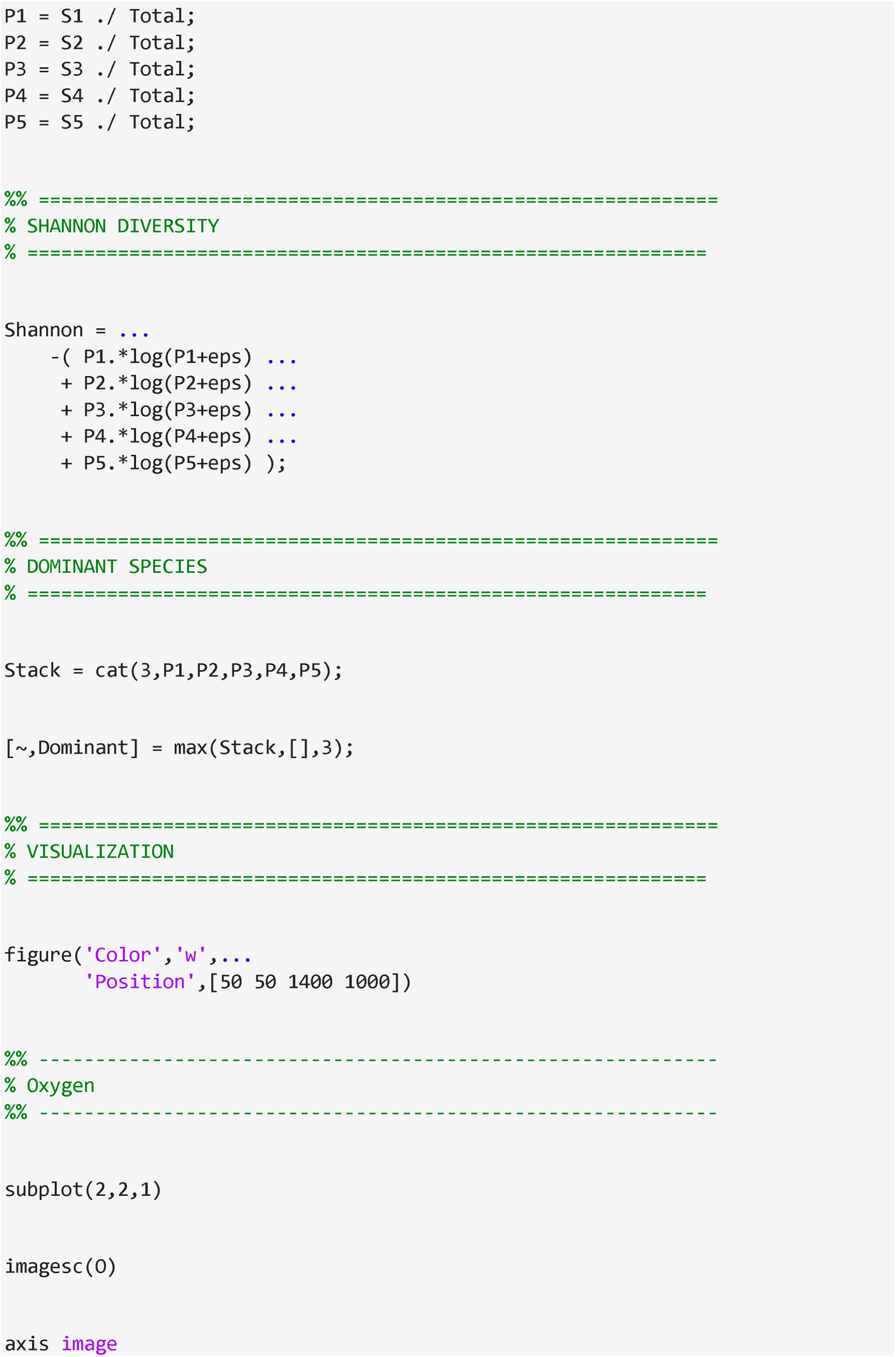
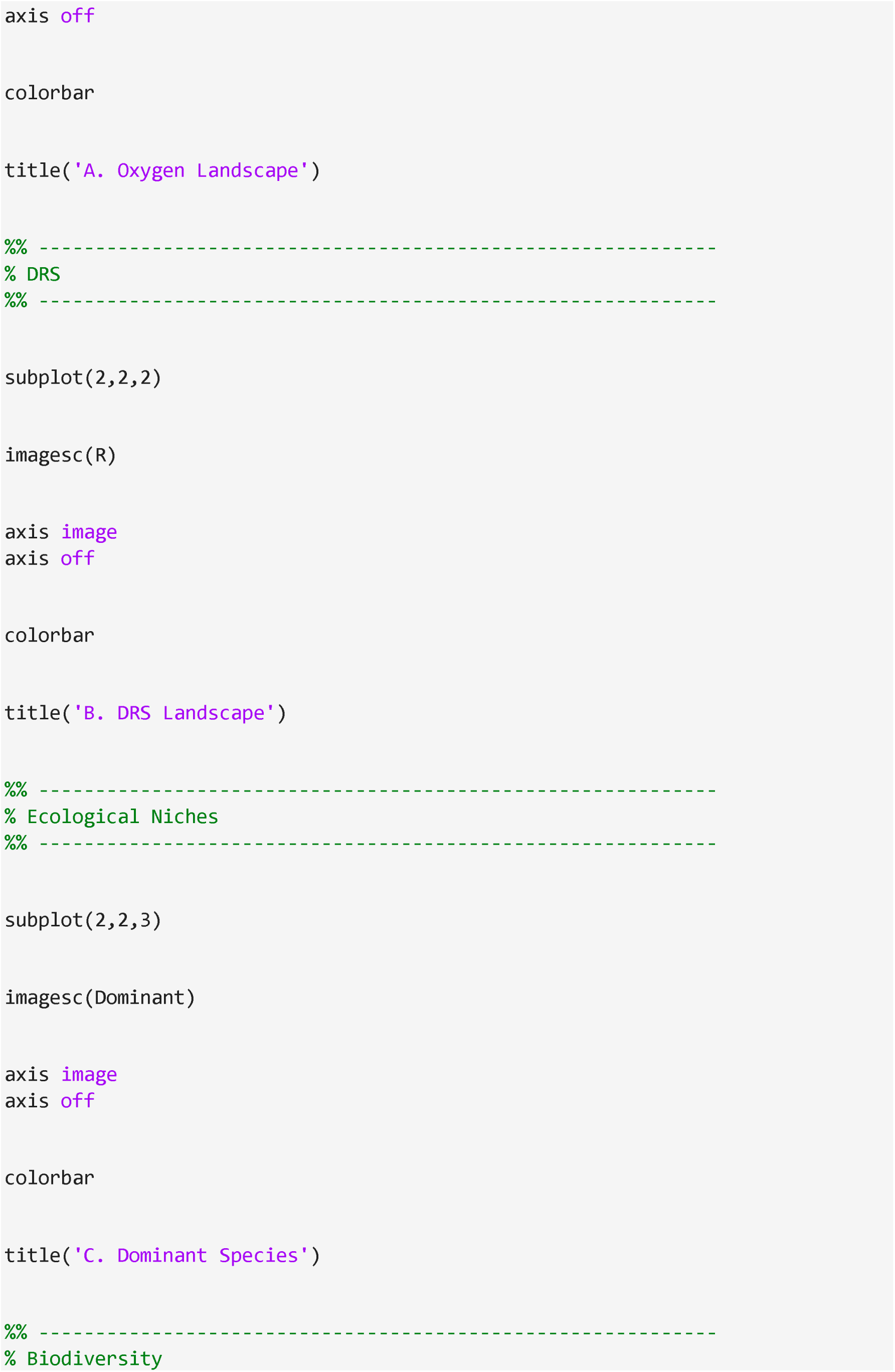
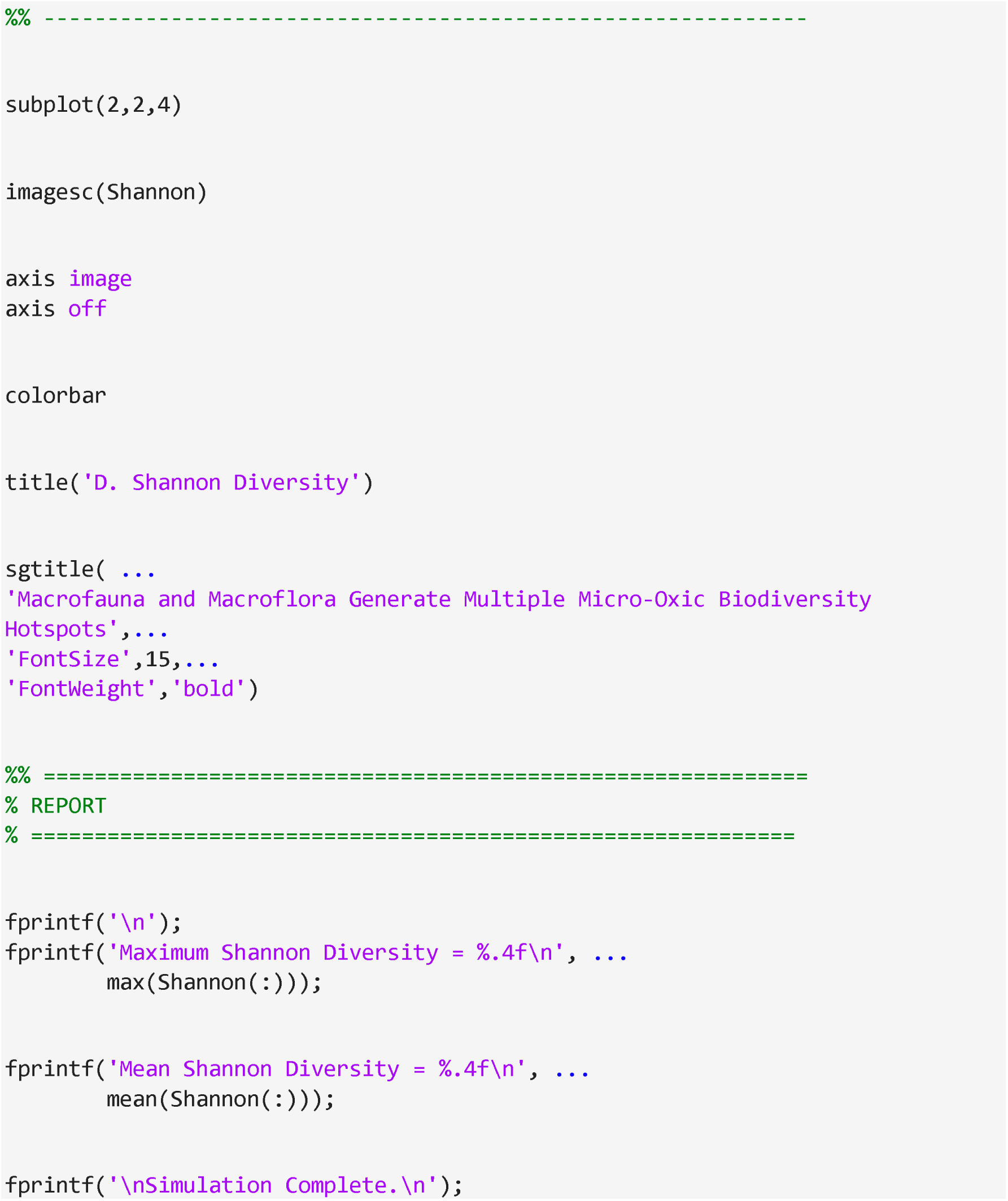

